# Spontaneous Dynamics Predict the Effects of Targeted Intervention in Hippocampal Neuronal Cultures

**DOI:** 10.1101/2025.04.29.651327

**Authors:** Elisa Tentori, George Kastellakis, Marta Maschietto, Alessandro Leparulo, Panayiota Poirazi, Davide Bernardi, Luca Mazzucato, Michele Allegra, Stefano Vassanelli

## Abstract

Microstimulation is a powerful tool for causal interrogation of neural circuits and for therapeutic neuromodulation. However, predicting network-level responses to focal perturbations remains a major challenge. To address this problem in a tractable manner, we combine experiments on networks of hippocampal neurons cultured on high-density multielectrode arrays with spiking network model simulations. To characterize spontaneous and stimulation-evoked network dynamics, we employ a combination of direct electrophysiological readouts and information-theoretic measures. We find that single-channel stimulation reliably evokes a small set of site-specific, stereotyped network activity patterns. Remarkably, effective connectivity inferred from spontaneous activity captures the spatial organization of perturbation responses, enabling reliable ranking of stimulation-evoked effects across the network. Our spiking network model reproduces these observations and reveals the interplay between short-term synaptic depression and distance-dependent excitatory and inhibitory connectivity in shaping both spontaneous and evoked interactions at different effective scales. Spontaneous activity involves local structural routes, while perturbation-evoked responses engage comparatively longer, polysynaptic pathways. By unifying *in silico* modeling with experimental measurements, this work links spontaneous network structure to stimulation-evoked dynamics, suggesting that spontaneous effective connectivity may serve as a tractable proxy for stimulation targeting in recurrent circuits, with potential implications for the rational design of neuromodulatory interventions.

## 1. Introduction

Since classical studies on motor perception [1–3], electrical stimulation with implanted microelectrodes (i.e., microstimulation) has been used to map primary sensory and motor areas [4, 5], to investigate the mechanisms underlying robustness of computations to external interference [6–9], learning [10–12], error coding [13–15], and the principles of neural control [16, 17]. In clinical settings, microstimulation is employed to improve selectivity and efficacy of Deep Brain Stimulation (DBS) interventions aimed at modulating dysfunctional neural circuits in Parkinson’s disease [18] and other brain disorders, including epilepsy [19]. However, the microstimulation-based perturbation approach *in vivo* encounters significant technical challenges. First, the difficulty in precisely targeting neuronal subpopulations and identifying high efficacy electrodes. Currently, stimulation sites are selected through laborious trial-and-error procedures, which are time-consuming and potentially invasive. Significant progress could be achieved by the ability to predict and model the effects of microstimulation solely based on recordings of spontaneous activity, prior to any perturbation. Second, the mechanistic principles that explain the propagation of perturbations from stimulated neurons across neuronal circuits remain elusive, causing an experimental and theoretical gap that hampers the prediction of microstimulation-evoked responses.

Here, we sought to bridge this gap in a tractable setting using dissociated neuronal cultures on High-Density Multi-Electrode Arrays (HD-MEAs). Although dissociation disrupts native anatomical organization, neuronal cultures self-organize into recurrent active circuits that preserve core cellular and synaptic mechanisms, including excitatory-inhibitory interactions and short-term synaptic plasticity [20, 21]. These networks generate robust spontaneous dynamics that organize into collective spatiotemporal patterns [22–25], whose richness is constrained by mesoscale network connectivity [26] and excitation-inhibition balance [20, 21]. Importantly, local cultured circuits can be investigated under controlled conditions in the absence of afferent projections mediating behavioral and cognitive modulations, allowing to isolate the effects of perturbations on circuit dynamics [27]. Combined with HD-MEA technology, cultured neural networks enable simultaneous largescale recording and flexible microstimulation [28, 29] under conditions that are difficult to achieve *in vivo*.

Our strategy combines three steps. First, we implement an experimental protocol to map the network-wide footprint of focal perturbations *in vitro*, by systematically stimulating a set of different sites and recording responses across the network. We estimate the Interventional Connectivity (IC) between stimulated and recorded electrodes [30] to obtain a perturbation map – or *perturbome* [31], i.e. the map of network responses to perturbations, reflecting pairwise, microstimulation-induced causal influences. Second, we test whether spontaneous activity alone predicts these perturbation effects. To this end, we estimate directed effective interactions from ensemble spike trains to reconstruct the circuit’s Effective Connectivity (EC) [32–34]. We find that EC captures the source-specific spatial organization of the IC perturbome beyond what is expected from geometric proximity alone, and predicts which stimulation sites exert the strongest network-wide effects. Third, to investigate candidate mechanisms underlying this correspondence, we develop a minimal biophysically grounded Spiking Neural Network (SNN) model with distance-dependent connectivity, realistic axonal delays, local inhibition, and presynaptic Short-Term Depression (STD) [35–38]. The model reproduces the main empirical features of both EC and IC. It reveals that EC primarily reflects direct synaptic connections, whereas IC engages multisynaptic pathways. In this framework, the model suggests that STD is required for sustained post-stimulation modulation, underlying the distributed spatial effects of perturbation encoded in the perturbome, while the spatial spread of perturbation responses is shaped by distance-dependent recurrent connectivity.

## 2. Results

### Terminology

Throughout the paper, the term *focal* refers to single-channel stimulation with simultaneous multi-unit recording. In the experimental sections, directed interactions are defined from stimulation channels (*sources*) to recording channels (*receivers*). To enable direct comparison with simulations, we use the same source-receiver notation in the model sections; *in silico* receiver neurons are also referred to as *postsynaptic*.

#### Focal perturbations reveal distance-dependent motifs *in vitro* and *in silico*

To test whether spontaneous directed interactions predict perturbation-evoked response profiles, we recorded dissociated rat hippocampal cultures with HD-MEAs (MaxOne, MaxWell Biosystems) using a two-phase protocol (Figure 1; Methods 4.1). From the 26,400 available electrodes, we selected a fixed map of *N* ≈ 1024 active channels centered on the most active regions of the culture; this map was kept constant throughout the experiment as a common spatial reference. In phase 1 (Figure 1a), we recorded 30 minutes of spontaneous spiking activity and estimated Effective Connectivity (EC) using delayed Transfer Entropy (TE). In phase 2 (Figure 1b), we systematically stimulated *M* = 20 selected electrodes, one site at a time, collecting 200 trials per site to map evoked responses (Methods 4.1). For each stimulation-recording pair (*i, j*), we computed Interventional Connectivity, *IC*_*i*→*j*_, as the Kolmogorov-Smirnov (KS) distance between pre- and post-stimulation spike-count distributions at receiver *j* (Methods 4.3).

**Figure 1:**
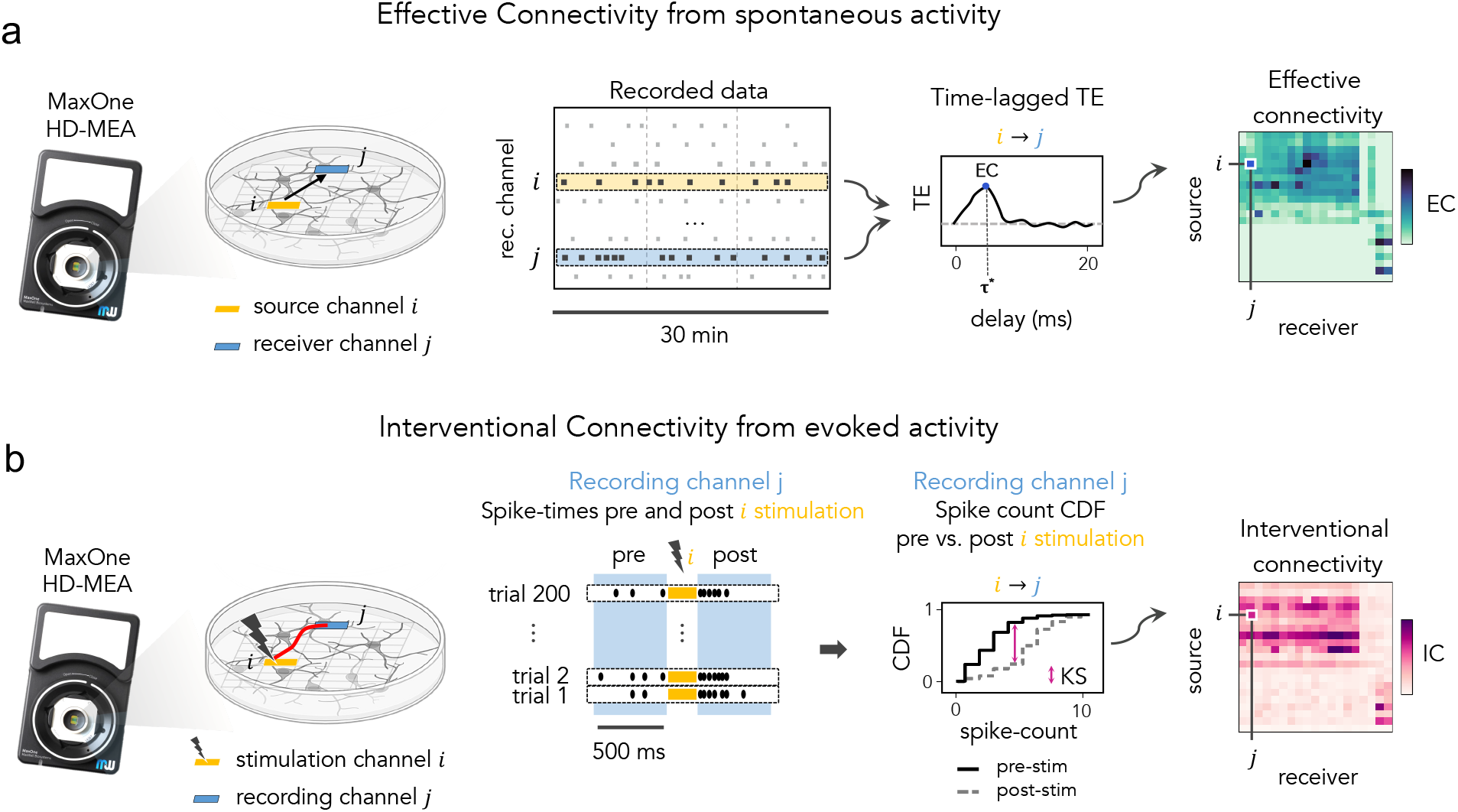
Experimental protocol for mapping spontaneous and perturbation-evoked directed interactions. A two-phase protocol design yields Effective Connectivity (EC) from spontaneous spiking activity and Interventional Connectivity (IC) from focal perturbations. **(a) Spontaneous activity phase: EC estimation**. A fixed map of *N* ≈ 1024 highly active channels is selected once at the beginning of the experiment and kept constant throughout. Spontaneous spiking activity from all *N* channels is recorded for 30 min. Spike trains are discretized into 0.3-ms bins and binarized. Then, delayed Transfer Entropy (TE) is computed over multiple time lags. For each ordered pair (*i, j*), the maximum TE across lags defines the directed EC weight. Repeating this computation over all pairs yields the full *N* × *N* EC matrix (right panel). **(b)** *Perturbation phase: IC estimation*. A subset of *M* channels is stimulated, one site at a time, while recording all *N* channels continuously. Each stimulation channel is tested across *N*_*T*_ = 200 trials, for a total of *M* × *N*_*T*_ = 4000 trials, with randomized stimulation order and a fixed 4-s inter-stimulation interval to allow recovery to baseline. For each stimulation-recording pair, IC is computed by comparing pre- and post-stimulation activity at the receiver.

The Results below are shown for two representative hippocampal cultures with distinct mesoscale organization: modular preparation with spatially segregated clusters (Culture 1; Figure 2a) and a spatially uniform preparation (Culture 2; Figure 2b). These empirical topologies also define reference architectures used for the corresponding *in silico* simulations.

**Figure 2:**
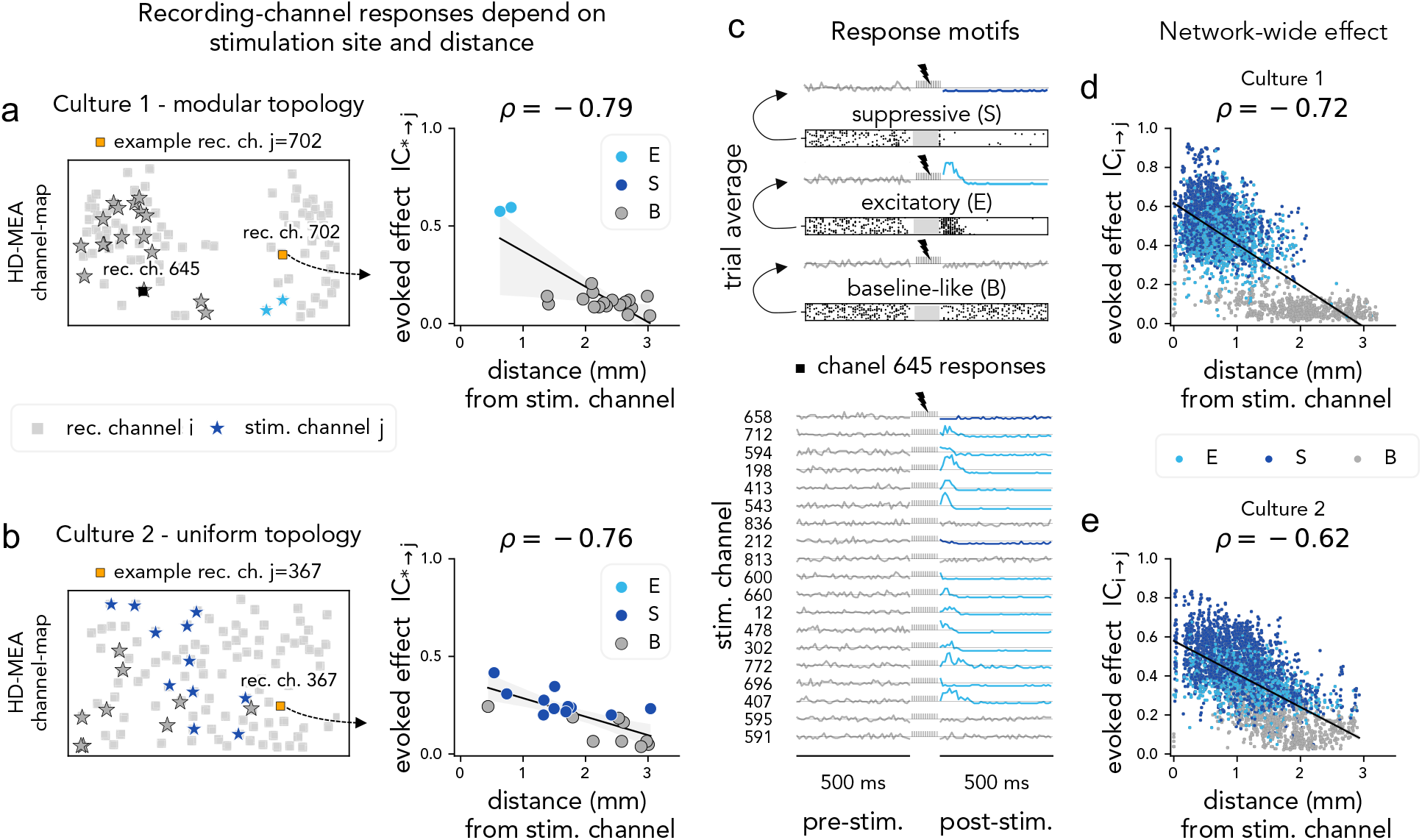
Stimulation-evoked responses are source-specific and distance-dependent. **(a**,**b)** For each culture, one example receiver channel is highlighted (orange square; *j* = 702 in culture 1, *j* = 367 in culture 2) on the HD-MEA map (gray squares). Stimulation channels (stars) are colored by the response motif evoked at that receiver (E, excitatory; S, suppressive; B, baseline-like). Right: evoked effect *IC*_*i*→*j*_ versus Euclidean distance from stimulation site *i*. The line shows the fitted monotonic trend, and *ρ* denotes Pearson correlation between *IC*_*i*→*j*_ and distance. **(c) Definition of response motifs from trial-averaged activity**. *Top:* representative S, E, and B motifs across *N*_*T*_ = 200 trials, shown as rasters (bottom) and trial-averaged firing rates in 10-ms bins (top). *Bottom*: trial-averaged responses of recording channel 645 (culture 1) to all stimulation channels (rows), aligned to stimulation onset; 500 ms pre- and post-stimulation windows. **(d**,**e)** Population-level distance dependence across stimulation-recording pairs (*i* → *j*) for the top 200 receivers ranked by IC in-strength, for culture 1 (d) and culture 2 (e). Each point is one pair, color-coded by motif; the line shows the fitted trend, and *ρ* is the Pearson correlation.

Focal perturbations elicited reproducible, source-specific effects. For a fixed receiver, changing the stimulated source produced distinct response motifs and effect magnitudes (Figure 2a,b): proximal stimulation typically yielded robust modulations, whereas distant sources more frequently resulted in baseline-like profiles. We classified trial-averaged responses into three stereotyped motifs (Figure 2c): baseline-like (B), suppressive (S), and transient excitation followed by suppression (E). Their consistency across repeated trials supports the robustness of the *IC*_*i*→*j*_ estimates for individual source-receiver pairs.

At the population level, IC magnitude decayed with source-receiver distance in both cultures (Figure 2d,e), but with distinct spatial profiles. In Culture 1 (modular), the decay was steeper and more spatially confined; in Culture 2 (uniform), it was smoother and closer to isotropic. To improve signal-to-noise ratio and visualization, population-level distance analyses were restricted to the top 200 receivers ranked by IC in-strength. Notably, a subset of long-distance pairs still showed sizable IC values (examples in Figure 2a,b, right), suggesting heterogeneous long-range propagation routes beyond the average spatial decay.

#### Response motifs and propagation profiles are reproduced in silico

To mechanistically interpret the relation between EC and IC, we applied the same two-phase protocol to a spatially embedded excitatory-inhibitory Spiking Neural Network (SNN) model (Methods 4.2). The model reproduces the two-dimensional layout of the cultures and includes distance-dependent excitatory coupling (Exponential Distance Rule; EDR), axonal delays, and local recurrent inhibition. Activity-dependent synaptic efficacy was modeled through presynaptic STD, enabling history-dependent responses to focal input. Using the same protocol as *in vitro*, we estimated EC from spontaneous activity and IC from focal perturbations. Because structural connectivity is known *in silico*, we could directly compare direct structural links with the broader interaction patterns recovered by EC and IC.

The model recapitulated the main empirical features of stimulation-evoked responses (Figure 3). Across stimulation trials, individual postsynaptic neurons displayed reproducible, source-dependent motifs matching the three classes observed *in vitro* (E, S, B; Figure 3a). In the model, these motifs arise from activity-dependent synaptic dynamics: stimulation transiently recruits excitatory and inhibitory pathways, and STD produces a post-stimulation modulation that persists beyond the stimulation train. Consistently, removing STD strongly reduced post-stimulation modulation under identical structural connectivity and stimulation parameters (Supplementary Figure S9), indicating that an adaptive synaptic mechanism is required for the protocol to yield diversified response motifs.

**Figure 3:**
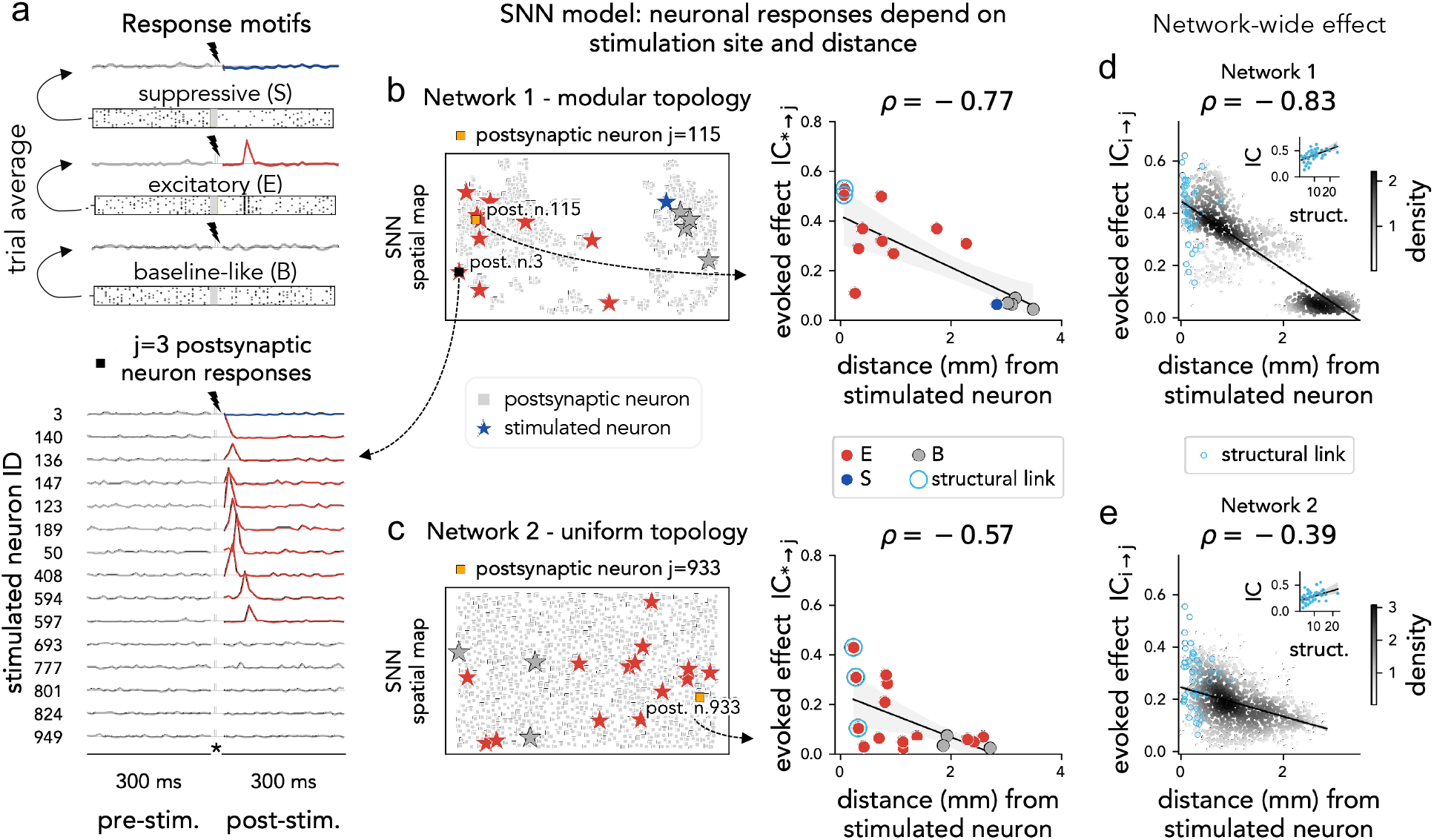
SNN model reproduces stimulation-evoked motifs and distance-dependent propagation. **(a) Response motifs defined from trial-averaged activity.** Top: examples of suppressive (S), excitatory (E), and baseline-like (B) motifs. Bottom: trial-averaged responses of one example postsynaptic neuron (*j* = 3) to stimulation from multiple sites (one row per stimulated neuron), aligned to stimulation onset. **(b) Modular network (Network 1)**. Left: spatial layout with stimulated neurons (stars) and an example postsynaptic neuron highlighted (orange square). Right: evoked effect *IC*_*i*→*j*_ for the selected postsynaptic neuron plotted against distance from each stimulated neuron; colors indicate response motif (E, S, B) and *ρ* is Pearson correlation. Cyan markers denote pairs with a direct structural connection. **(c) Uniform network (Network 2)**. Same conventions as in (a). **(d**,**e)** Population-level distance dependence of IC across the top *K* = 100 most responsive postsynaptic neurons (ranked by IC in-strength); point density is shown by grayscale shading and the line indicates the fitted trend. Cyan markers denote pairs with a direct structural connection. *Insets:* scatter of *IC*_*i*→*j*_ versus structural weight for directly connected pairs (cyan), showing a positive association between direct-link strength and evoked effect.

While STD accounts for the temporal structure of the motifs, their spatial organization is set by the spatially embedded connectivity architecture. In both the modular and uniform networks, the evoked effect decreased with distance from the stimulated neuron, and baseline-like responses predominated at longer distances (Figure 3b, 3c). This dependence differed across architectures: in the modular network, long-range effects were attenuated because the two modules are only sparsely connected and long connections are predominantly cross-module ones; in the uniform network, effects spread more isotropically (Figure 3b-e). In a control analysis, we simulated a model variant without EDR, i.e. distance-independent excitatory wiring with fixed node positions. Motif-like temporal responses persisted, but IC lost its distance-dependent footprint (Supplementary Figure S7, Figure S8).

Because ground-truth connectivity is accessible in the model, we tested how stimulation-evoked influence relates to the underlying wiring (Figure 3d, 3e). To improve visualization, we restricted the analysis on the top 100 most responsive targets, as ranked by IC in-strength; the corresponding all-pairs analysis produces fully consistent, albeit less sharp results. Within directly connected pairs, *IC*_*i*→*j*_ showed substantial variability, yet increased with structural weight, indicating a graded contribution of direct coupling strength to perturbation impact (Figure 3d, 3e, insets). At the same time, non-zero *IC*_*i*→*j*_ values also occurred for pairs lacking a direct structural connection, showing that focal perturbations propagate beyond direct links through indirect recurrent pathways, as we examine in more detail below.

#### Spontaneous EC Predicts Stimulation-evoked IC *in vitro*

We next asked whether effective interactions inferred from spontaneous activity predict the “perturbome” evoked by focal microstimulation (Figure 4).

**Figure 4:**
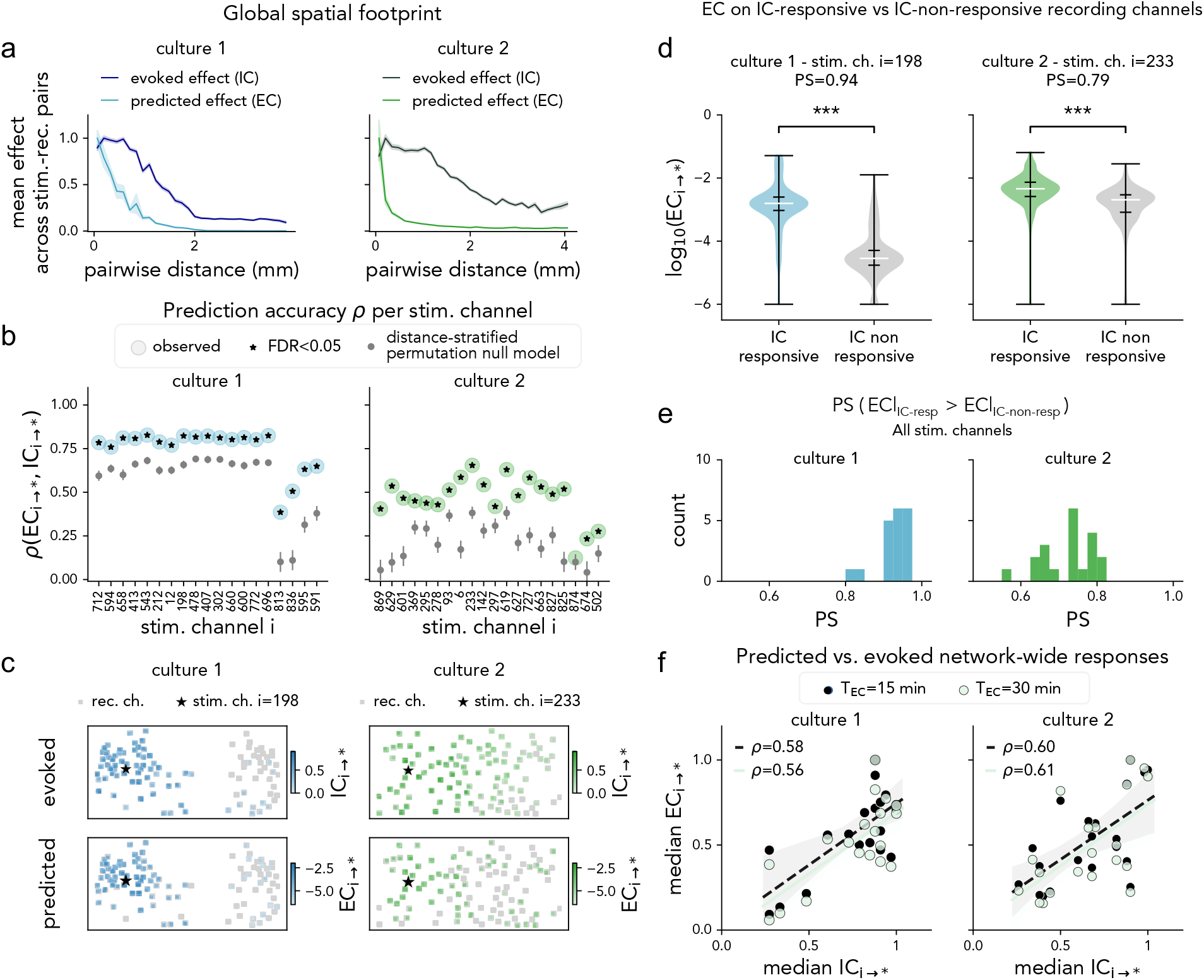
*In vitro* prediction of stimulation-evoked perturbation patterns from spontaneous effective connectivity. **(a) Global spatial footprint**. Mean evoked effect (IC) and mean predicted effect (EC) across stimulation-recording pairs are shown as a function of pairwise distance (culture 1 and culture 2). Both decay with distance, with a steeper decay for EC than IC. **(b) Source-wise prediction beyond distance**. For each stimulation channel *i*, prediction accuracy was computed as Spearman *ρ*(*EC*_*i*→∗_, *IC*_*i*→∗_) on raw outgoing profiles (before significance thresholding). Observed *ρ* was compared against a distance-stratified permutation null (15 distance bins; *N*_perm_ = 5,000 row-wise shuffles preserving within-bin IC distributions). Gray dots represent the null distribution; stars denote channels significant after Benjamini–Hochberg FDR correction across sources (*α* = 0.05; Methods). **(c) Example source patterns**. Representative source-specific maps for each culture showing evoked and predicted outgoing patterns on the HD-MEA layout, using significance-thresholded EC and IC. Significant receivers are colored; non-significant receivers are shown in gray. **(d) Edge-level separability (examples)**. Distributions of outgoing EC weights for the example sources in (c), partitioned by IC-significance (IC-responsive vs. IC-non-responsive receivers). Asterisks indicate one-sided permutation tests on the median difference Δ = median(*EC*_resp_) − median(*EC*_non-resp_) (*N*_perm_ = 20,000, group sizes preserved; Methods). **(e) Edge-level separability across sources**. Source-wise separability summary reported as probability of superiority (PS, from Cliff’s delta) for *EC*|*IC*_resp_ versus *EC*|*IC*_non-resp_; significance from the same one-sided median-permutation framework as in (d), with 95% bootstrap CI on Δ (*N*_boot_ = 5,000 resamples; Methods). **(f) Predicted vs evoked network-wide responses**. Source-level comparison of median outgoing *EC*_*i*→∗_ versus median outgoing *IC*_*i*→∗_ across stimulation channels, for EC estimated from 15-min and 30-min spontaneous recordings, showing robustness of source ranking across recording durations.

In both modular and uniform cultures, predicted (EC) and evoked (IC) effects decayed with stimulation-receiver distance (Figure 4a), but with different spatial extent: IC showed a broader footprint, while EC remained more localized around the stimulation source. This divergence suggests that stimulation may recruit routes that are only weakly expressed during spontaneous activity – a point that we test mechanistically in the model section below.

Quantitatively, the correlation between predicted and evoked patterns, *ρ*(*EC*_*i*→∗_, *IC*_*i*→∗_), averaged 0 75 ± 0 11 across stimulation channels in culture 1 and 0.54±0.13 in culture 2. Because both measures covary with distance, their strong correlation may be entirely due to this confounder. To test this, we used a null model where EC strengths were shuffled among edges with matching between-node distance (Figure 4b). Distance-matched surrogates yielded markedly lower correlations (*ρ*_null model_ = 0.56 ± 0.19 in culture 1; *ρ*_null model_ = 0.21 ± 0.10 in culture 2), indicating that EC captures perturbation effects beyond geometric proximity (Figure 4b,c).

We complemented source-wise correlation with an edge-level separability analysis, to test whether EC distinguishes stimulation-responsive receivers from receivers that remain at baseline. For each source, outgoing EC edges were labelled as “responsive” or “non-responsive” depending on whether IC was significant, and separability was quantified as the difference between the median EC values of the two groups. EC medians were systematically higher for responsive than non-responsive edges (Figure 4d). This separation, quantified as Probability of Superiority (PS) for each source, was more pronounced in the modular culture than in the uniform one (PS = 0.93 ± 0.04 vs 0.72 ± 0.06; Figure 4e), likely reflecting the stronger structural constraints imposed by modularity. The inverse conditioning analysis (IC grouped by EC significance) yielded similar results (Supplementary Figure S5).

Finally, we evaluated the operational utility of our framework by testing whether spontaneous EC could effectively rank stimulation sites by their global network impact. We summarized the outgoing influence of each source as the median effect across the full receiver population in both cultures. The source ranking was well preserved and remained stable even when EC was estimated from only 15 min of spontaneous activity (Figure 4f). This indicates that brief spontaneous recordings can provide a practical proxy for prioritizing high-impact stimulation channels, potentially bypassing the need for exhaustive trial-and-error screening in online-calibrated experimental protocols.

#### Model-based mechanism of EC–IC correspondence

To interpret the experimental EC-IC correspondence mechanistically, we applied the same analysis pipeline to the SNN model, where ground-truth structural connectivity is known and stimulation-driven propagation can be tracked directly.

##### The model reproduces in vitro EC–IC spatial organization and source-wise predictability

The SNN model recapitulated the geometric signatures observed *in vitro*: both EC and IC decayed with source–target distance, yet IC showed a systematically broader footprint than EC (Figure 5a). Both functional readouts extended beyond direct structural connections, with EC remaining near-field biased and IC showing the broadest spatial spread. Thus, the *in silico* results reproduced the ordering seen *in vitro*, with IC extending farther than EC. Despite this footprint mismatch, source-wise prediction from EC to IC remained robust. For each stimulated source *i*, we computed the Spearman correlation between outgoing profiles, *ρ*(*EC*_*i*→∗_, *IC*_*i*→∗_). Mean accuracy was *ρ* = 0.60 ± 0.09 in Network 1 (modular) and *ρ* = 0.53 ± 0.10 in Network 2 (uniform), and exceeded the corresponding distance-matched null distributions (*ρ*_null model_ = 0.31±0.12 and 0.09 ± 0.06, respectively; Figure 5b). This indicates that EC carries source-specific information beyond geometric proximity, consistent with the *in vitro* result.

**Figure 5:**
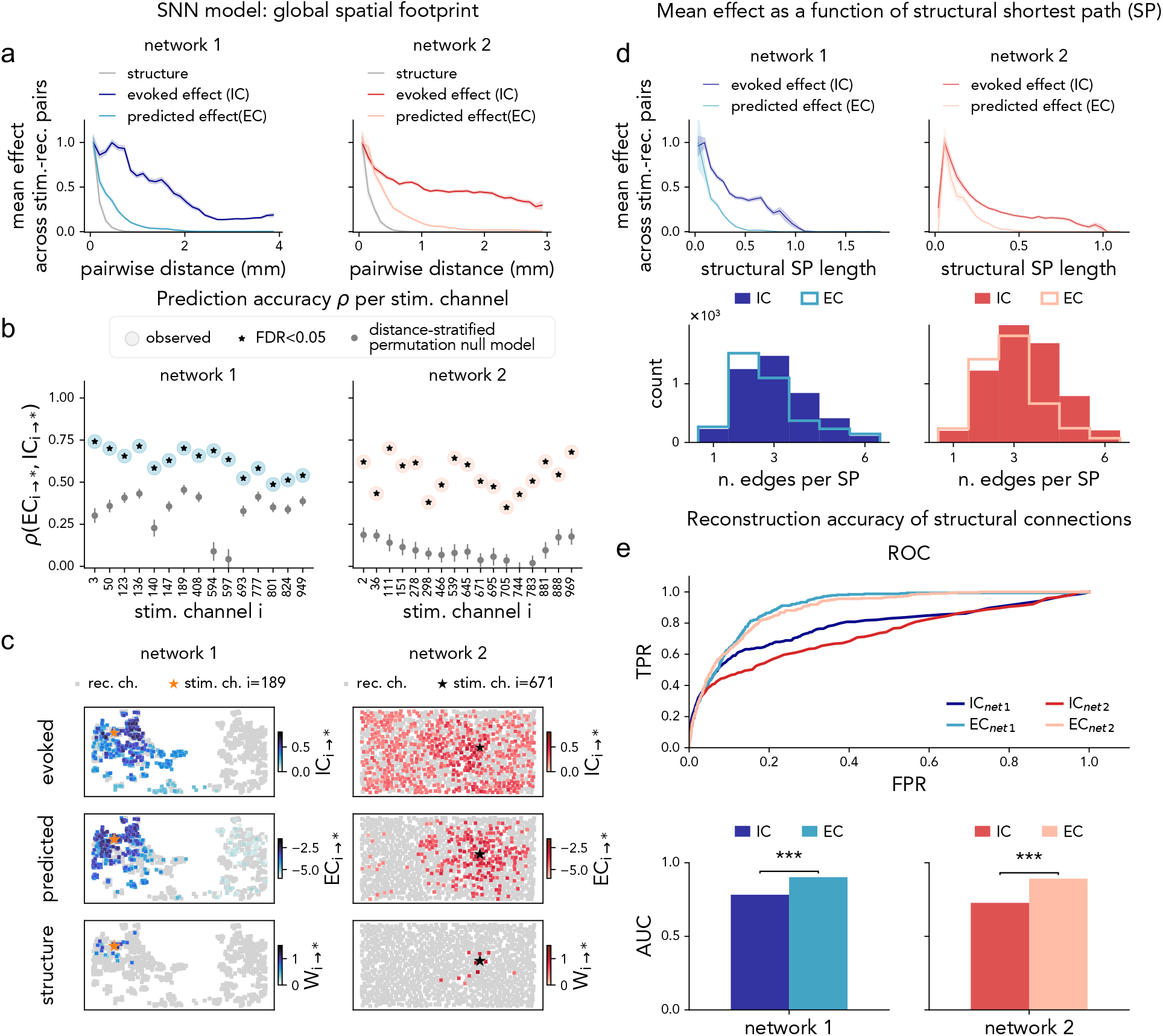
Spiking model reveals that spontaneous EC reflects the structural core, whereas IC recruits broader polysynaptic pathways. **(a) Global spatial footprints *in silico***. Mean structural weight, predicted effect (EC), and evoked effect (IC) are shown as a function of Euclidean distance for the modular (Network 1) and uniform (Network 2) topologies. EC defines a near-field functional footprint that extends farther than the mean structural weight profile, whereas IC shows the broadest spatial spread, reaching distal targets through recurrent propagation. **(b) Prediction accuracy of focal perturbations**. For each stimulated neuron *i*, accuracy was quantified as Spearman *ρ*(*EC*_*i*→∗_, *IC*_*i*→∗_) computed on raw outgoing profiles and compared with a distance-stratified permutation null (Methods). Stars denote significance after FDR correction. **(c) Topological vs. functional mapping**. Representative examples of structural outgoing weights (*W*_*i*→∗_), predicted EC patterns (*EC*_*i*→∗_), and evoked IC responses (*IC*_*i*→∗_) for fixed stimulation nodes. IC patterns recruit broader network regions than those directly connected to the source or captured by spontaneous EC. **(d) Polysynaptic organization of causal flow**. Mean IC and EC are plotted against SP length between source-receiver pairs. IC remains elevated over longer structural paths, whereas EC is concentrated on shorter paths. Bottom: corresponding distributions of the number of edges contributing to each SP bin. **(e) Recovery of direct structural links**. ROC curves and AUC for EC and IC in classifying direct structural connections. EC shows higher discriminability than IC in both networks (*p* < 0.001), indicating tighter alignment with direct structural wiring.

Representative source maps (Figure 5c) further show that, although EC and IC are not identical at the edge level, they remain topographically aligned: both spread over the structural neighborhood of the source, with IC extending farther than EC.

##### Shortest-path and ROC analyses show that EC and IC sample different structural routes

To explain why EC predicts IC despite its narrower spatial footprint, we quantified structural shortest paths (SPs) between all node pairs (Methods 4.4; Figure 5d). Mean non-zero edge weights decreased with SP length for both metrics, but more steeply for EC (Figure 5d, top), indicating that EC preferentially emphasizes shorter/stronger structural routes, whereas IC remains comparatively elevated on longer/weaker routes. We next computed, for each detected edge (*i, j*), the number of hops in the structural shortest path linking *i* and *j*, and compared the hop-count distributions of EC-detected and IC-detected edges (Figure 5d, bottom; non-normalized counts). IC showed higher total counts than EC, indicating broader edge recruitment under perturbation and sparser spontaneous detection by EC. In both networks, EC was left-shifted relative to IC (in Network 1, the modal hop count was 2 for EC versus 3 for IC), consistent with preferential sampling of shorter structural routes.

We then used a ROC analysis to quantify how accurately each functional measure recovered direct structural edges. Both EC and IC classified direct structural links above chance, but EC yielded higher AUC in both topologies (Figure 5e). This pattern is consistent with tighter alignment of EC to direct wiring, whereas IC is more strongly shaped by indirect propagation.

These results provide a mechanistic account of the experimental findings: EC and IC sample the same underlying architecture at different effective scales, with EC weighted toward shorter structural routes and IC extending to longer-range, polysynaptic interactions.

## 3. Discussion

Microstimulation has a huge potential for probing neuronal circuits and therapeutic applications. However, this potential cannot be fully realized without a stronger control of the spatiotemporal effects of stimulations within neuronal networks, and a more solid understanding of the underlying mechanisms. Due to their experimental accessibility, neuronal cultures plated on HD-MEAs are a natural testing ground for the study and development of neurostimulation paradigms, and they have been extensively used in previous studies - e.g., to modulate synchronization in a neural population [38–42], sculpt network connectivity to enable specific computation [43–48], or test novel multi-site stimulation protocols [49–51]. Here, we combined an innovative experimental design and computational modeling to obtain a mechanistically grounded model consistent with the main experimental observations of the effects of microstimulation on cultured neuronal networks.

A major pillar of the present work is the employment of a systematic experimental protocol whereby single sites are stimulated in succession and the evoked responses are monitored throughout the network. While this design was previously tested in *vivo* [30, 52], we demonstrated its feasibility and robustness *in vitro*, showing that perturbation-induced effects were reproducible and exhibited well-defined spatial specificity. The repertoire of network’s responses comprised three stereotyped firing pattern motifs: baseline-like, transient excitatory followed by suppression, and fully suppressive. By stimulating multiple channels, we achieved a reproducible characterization of the so-called network’s *perturbome* [31], i.e., the map of pairwise perturbation effects induced by stimulation in the network.

Our finding that spontaneous activity carries predictive information about perturbation spread in the network has practical implications for *in vitro* and, potentially, *in vivo* applications of microstimulation. The selection of stimulation sites is a central bottleneck in the design of neuromodulatory protocols [53], and we showed that Effective Connectivity (EC) calculated from spontaneous activity reliably ranked stimulation sites by their network-wide impact. Notably, EC can be computed rapidly and with limited data, providing a reliable and fast proxy that could bypass the need for trial-and-error exploration and favor *online* identification of effective stimulation targets at low computational cost. Our paradigm is of potential applicability to *in vitro* neuronal preparations, including not only classical neuronal cultures, but also brain organoids, which are increasingly being studied through multi-electrode arrays [54–56]. Although some previous works investigated the link between spontaneous activity and stimulation in cultures, their focus was mainly restricted to collective events (bursts or global activity states) [41, 48, 57]. In contrast, our results show that EC reproduced several spatial features of the perturbome, including source-specific footprint and separability between responsive and non-responsive channels.

By developing a reliable SNN-based computational model of *in vitro* neural activity, we obtained deep insight into the culture’s dynamics and the mechanisms supporting the propagation of stimulation effects. This model could be used in later studies to investigate specific theoretical hypotheses about the dynamics of neural networks (e.g., excitation-inhibition balance or linearity [12]) and for *in silico* exploration of advanced neurostimulation protocols. Within this modeling framework, reproducing the empirical response profiles and motifs required two key ingredients: i) presynaptic STD and ii) spatially embedded, distance-dependent excitatory-inhibitory recurrent connectivity with axonal delays. STD was required to produce lasting reverberation after stimulation: for a fixed structural connectivity and E/I organization, removing STD resulted in short-lasting, spatially confined peak responses. This finding is consistent with the presence of short-term adaptation mechanisms in neuronal cultures, as supported by several experiments [41, 58] and previous modeling efforts [38]. The empirical spatial footprint of responses was reproduced by using an exponential-distance-rule (EDR) synaptic connectivity. While EDR is widely observed across cortical networks in both rodents and primates [59, 60], its validity in dissociated cultures was suggested by previous experimental and modeling work [33, 34, 61] and our findings support the plausibility of this hypothesis. Further studies could leverage cutting-edge techniques to directly measure synaptic strength and yield a clear-cut test of this hypothesis [62–65]. Our model provides a clarifying picture of the mechanisms underlying the observed response phenomenology. In the model, the existence of stereotyped response motifs was shown to depend on a delicate interplay between excitation and inhibition: the recruitment of excitatory neurons near to the stimulation site produces an intial excitatory response, whereas delayed inhibitory feedback eventually produces suppression, with the balance between the two determining whether the net effect is transiently excitatory or fully suppressive. This picture aligns with, and supports, conclusions of previous computational works trying to explain why stimulation can cause a local amplification followed by inhibition [66, 67], which highlighted the interplay of circuit structure (characterized by feed-forward excitatory and recurrent inhibitory connections) and short-term plasticity. The model also allows us to relate IC and EC to the underlying circuit wiring. EC-IC alignment thus reflects a shared dependence on the recurrent spatial backbone: EC samples shorter and stronger pathways, whereas IC extends along weaker, multi-step routes. This explains the strong node-level EC-IC correspondence observed *in vitro* and clarifies why EC predicts which sites to stimulate but not the full extent of the reverberation triggered by stimulation. Obviously, the present work has several limitations. First, our reconstruction of the perturbome was partial, as we used a limited number (20) stimulation channels. However, this choice was instrumental to characterize the response stability: since the latter has been established, future experiments could easily increment the number of stimulation targets (e.g., up to 100) while shortening the number of trials per channel, allowing for a spatially finer perturbome reconstruction. Secondly, besides the recruitment of different pathways, the partial alignment of EC and IC may also depend on our usage of a relatively simple metric to characterize EC (the transfer entropy). Future work may explore more advanced causal analysis tools (such as convergent cross mapping [30, 68]) in search of an even better prediction of stimulation responses from spontaneous activity. Finally, while our computational model can qualitatively reproduce all the main features of the observed spontaneous and stimulation-evoked activity, model refinements, increasing the model’s biological plausibility, are required for a better quantitative match. In particular, responses to stimulation in the model tend to be more transient than in experimental cultures. Future work should address this gap, e.g., by introducing a slow mechanism in the model (such as, e.g., a slow NMDA-related synaptic plasticity).

In conclusion, by showing that spontaneous dynamics carry actionable information about how perturbations propagate, this work establishes a principled route to stimulation targeting: stimulation-site prioritization can be informed before intervention. Dissociated networks therefore become not only a tractable experimental system, but a testbed where predictive perturbation strategies can be developed, validated, and generalized to increasingly complex *in vitro* models.

## 4. Materials and Methods

### 4.1 Experimental and Modeling Framework

#### HD-MEA Setup

Experiments were performed using MaxOne High-Density Multi-Electrode Arrays (HD-MEAs; Maxwell Biosystems, Switzerland; https://www.mxwbio.com/products/maxone-mea-system-microelectrode-array/). The CMOS-based MaxOne platform comprises 26,400 platinum electrodes and allows simultaneous extracellular recording from up to 1,024 channels. Up to 32 of these can be dynamically connected to the stimulation module (DACs) for extracellular electrical input; in our experiments, 20 channels were assigned for stimulation. Each chip covers a 3.85×2.10 mm^2^ sensing area, providing an electrode density of 3,625/mm^2^ and 17.5 *Μ*m pitch. The chip is housed in a culture chamber integrated into a portable Recording Unit (RU) compatible with standard humidified incubators. The RU connects via a hub to a computer running the MaxOne Server, which interfaces with the licensed *MaxLab Live Scope V24*.*2* software. This software provides a graphical interface for control and visualization and a Python-accessible Application Programming Interface (API) for customized workflows. We operated in open-loop mode, integrating *MaxLab Live Scope V24*.*2* with custom Python modules for stimulation, acquisition, and spike detection.

#### HD-MEA Coating

The chip is integrated into the base of a circular chamber (19 mm internal diameter, 8 mm height) designed to contain the culture medium. Few days before neuronal seeding, microchips were sterilized in 70% ethanol (v/v) for 45 minutes, rinsed with sterile deionized water and air-dried under a sterile laminar hood. Then, they were pre-incubated at 37°C in L15 medium (see paragraph 4.1). On plating day, chips were rinsed and coated with 20 *μ*g/ml poly-L-lysine (PLL, Merck) for 2-h at room temperature, then dried before cell seeding.

#### Cell Cultures and Neuronal Plating

Primary hippocampal cultures were prepared from Wistar rat embryos (E18–E20) following a protocol adapted from [69]. Briefly, hippocampi were dissected from the brain under a sterile laminar hood, incubated with 0.08% Trypsin-EDTA, then dissociated via gentle trituration. After a preplating step to reduce the amount of glial cells, 120, 000 neurons/cm^2^ were seeded on PLL-coated chips in 100 *μ*l of L15 medium supplemented with 5% fetal bovine serum and 1% penicillin-streptomycin. After a 90-minute incubation at 37 °C in a 5% CO_2_ atmosphere, neurobasal medium supplemented with 1% Glutamax-1 and 2% B27 was added to fill the chamber. Two-thirds of the medium were refreshed every 3-4 days. All media and reagents were from Gibco (Thermofisher Scientific).

#### Ethics Statement

Wistar rats (Charles River) were housed under standard environmental conditions at the Animal Research Facility of the Department of Biomedical Sciences, University of Padua, in compliance with Italian animal welfare regulations (authorization numbers 522/2018-PR and 709/2023-PR).

#### Experimental Data and Spike Detection

Experiments were performed on dissociated hippocampal cultures between Days in Vitro (DIV) 23 − 33. During acquisition, chips were maintained in a humidified incubator at 37°C and 5% CO_2_. Neural activity was sampled at 20 kHz, and processed in real time through hardware-level filtering and threshold-based spike detection. Signals were high-pass filtered at 300 Hz, and spikes were detected at a threshold of five times the root-mean-square noise level above baseline, according to manufacturer guidelines. Connectivity estimation was based directly on these online-detected spike trains, allowing near real-time analysis without offline spike sorting.

#### Channel Map Selection

At the beginning of each experiment, a fixed recording map was defined to cover the most active regions of the culture. First, activity across all 26,400 electrodes was scanned using the *Activity Scan Assay* in *MaxLab Live Scope V24*.*2*, which ranks electrodes by spike amplitude. Then we chose the channel map, selecting the *N* ≈ 1,024 electrodes showing the highest spiking activity, constraining the spatial arrangement in 3 ×3 clusters of adjacent sites to optimize local sampling and maintain compatibility with optional spike sorting (not used here). The resulting *N*-channel recording map was kept fixed for the entire experimental session.

#### Experimental Design

Each experiment consisted of two consecutive phases: (i) spontaneous activity recording (30 minute session), used for causal flow inference (Effective Connectivity - EC), and (ii) focal electrical stimulation (4 h and 30 minute session), to quantify intervention-based causality (Interventional Connectivity - IC). The same culture and recording map were maintained throughout.

##### Spontaneous activity phase

Following channel map selection, spontaneous spiking activity was recorded for *T*_rec_ = 30 minutes from the *N* active electrodes identified by the Activity Scan Assay. Spike times were detected online as described above and used to compute EC across channel pairs (Figure 1a; see Section 4.3 for details). No external inputs were delivered during this phase.

##### Focal stimulation phase

In the second phase, we systematically probed network responses to focal perturbations, alternating single-channel electrical stimulation and multi-unit recordings. A subset of *M* = 20 stimulation channels was randomly selected from the fixed recording map. Stimulations were applied sequentially, one channel at a time, while activity from all *N* channels was recorded simultaneously. At each iteration, one stimulation channel was randomly chosen and received a train of biphasic electrical pulses [70] (amplitude 300 mV, frequency 200 Hz, duration 120 ms). Neural activity was recorded continuously to capture both baseline and evoked response. A fixed interstimulation interval of *T*_inter-stim_ = 4 s allowed the network to return to baseline before the next iteration (Figure S1a; Supplementary Information 4.4). This procedure was repeated sequentially across all stimulation channels and iterated for *N*_*T*_ = 200 trials per channel, ensuring full coverage of the stimulation set.

##### Experimental stability

Stimulation trials were randomized and interleaved across the *M* selected channels, to minimize long-term drift or stimulation-induced plasticity, preventing repeated activation of the same site within short intervals. Control analyses confirmed stable firing rate and burst statistics across the entire stimulation phase (Supplementary Information S6).

### 4.2 Microscopic Model

To replicate and mechanistically interpret the observed dynamics, and to validate our causality–inference protocol, we implemented an *in silico* Spiking Neural Network (SNN) model reproducing both spontaneous and stimulation-evoked activity, under the same protocol used for experiments. We generated a spatially embedded Excitatory-Inhibitory (E-I) network, driving spiking intrinsically bursting dynamics with an Izhikevich model [71, 72]. Communication delays and presynaptic Short-Term Depression (STD) [35– 38] were included to ensure realistic burst refractoriness and sensitivity to external perturbations. The network was designed to capture key structural and dynamical properties of dissociated hippocampal cultures, including sparse modular organization, recurrent excitation and inhibition, and activity-dependent depression of synaptic efficacy.

#### Network Architecture

We built spatially embedded E–I networks composed of *N*_SNN_ = 1000 neurons, with an E/I ratio of 4 : 1. E population was divided into Regular Spiking (RS) and Intrinsically Bursting (IB) neurons subsets, and I into Fast Spiking (FS) and Low Threshold Spiking (LTS) subsets, matching the most observed firing rate profiles observed in hippocampal cultures. Neurons were positioned in a two-dimensional domain, preserving minimum inter-node distance to prevent overlap. Two architectures were tested: a modular network with two spatially segregated clusters, characterized by dense intra-cluster and sparse inter-cluster connectivity, and a non-modular network in which neurons were uniformly distributed. Directed, sign-constrained connections (*E* → {*E, I*} > 0, *I* → {*E, I*} < 0) were assigned according to distance-based rules, differing by neuron type. Excitatory projections followed a broad Exponential-Distance Rule (EDR), 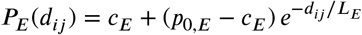, allowing long-range coupling across adjacent clusters. Inhibitory projections were local and range-limited within a radius *r*_*IE*_, *P*_*I*→*E*_(*d*_*ij*_) = *p*_*I*→*E*_ **1**[*d*_*ij*_ ≤ *r*_*IE*_]. Connection weights decayed exponentially with distance and were drawn from sign-truncated Gaussian distributions. Excitatory and inhibitory delays were distance-dependent.This spatial embedding defined a biologically grounded mapping between position and connectivity, enabling direct comparison of perturbation responses in modular and uniform networks. Parameters are listed in Supplementary Table 1.

#### SNN Model

Neuronal dynamics were modeled using Izhikevich neurons [71, 72] with presynaptic STD [35–38]. The single-neuron dynamics followed the coupled differential equations

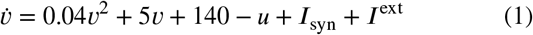

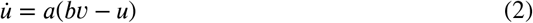

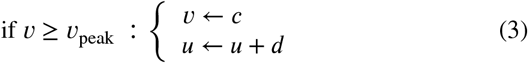

where *v* is the membrane potential and *u* the membrane recovery. Class-specific parameters (*a, b, c, d*) reproduced RS, IB, FS, and LTS firing behaviors. When spike was emitted, an absolute refractory period imposed to 3 ms for excitatory, 1 ms for inhibitory. Excitatory and inhibitory synaptic inputs were modeled as exponentially relaxing AMPA and GABA currents with delayed presynaptic impulses [38, 73]: 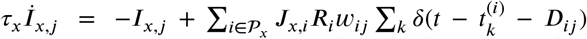, where *x* ∈ {AMPA, GABA^*x*^}, 𝒫_*x*_ denotes the corresponding presynaptic population (E or I)^*x*^, and *D*_*ij*_ are axonal delays (1–10 ms for excitatory, 1 ms for inhib^*ij*^itory). Presynaptic gains *J*_*x,i*_ were drawn from lognormal distributions (means: 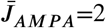 and 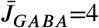) to capture release heterogeneity [37]. Each presynaptic neuron carried a resource variable *R*_*i*_ ∈ [0, 1] evolving under the depression-only limit of the Tsodyks–Markram model [35, 36]: 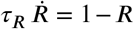, with *R* ← *β*_*E*/*I*_ *R* at each spike. This formulation implements depletion and recovery of neurotransmitter resources, with *β*_*E*_ = 0.1, *β*_*I*_ = 0.8, *τ*_AMPA_ = 5 ms, *τ*_GABA_ = 20 ms, and *τ*_*R*_ = 0.5 s [37, 74]. Note that the inhibitory term here combines the effects of both fast GABA_*A*_ and slow GABA_*B*_ receptors, which are known to contribute to the local response to microstimulation [75]. All neurons received an independent excitatory shot-noise drive *I*^ext^ (*t*) = *A*_*E*_Σ_*k*_ *δ* (*t* − *t*_*k*_), where *t*_*k*_ were drawn from a Poisson process of rate *ν*_*E*_, providing a stationary excitatory background that represents intrinsically firing neurons - essential for sustaining network activity and known to exist in neuronal cultures [76, 77]-as well as unmodeled noise sources. This combination of adaptive spiking, distance-dependent delays, and synaptic resource depletion reproduced the alternation of network bursts and quiescent periods observed experimentally. Full parameter values are reported in Supplementary Table 2.

#### In silico Experimental Design

Simulations used a fixed time step Δ*t*_SNN_ = 1 ms. We reproduced the same two-phase protocol applied *in vitro*: (i) a spontaneous activity block (*T*_rec, SNN_ = 30 min), and (ii) a focal stimulation block implementing sequential single-site perturbations. For stimulation, we randomly selected *M*_SNN_ = 15 neurons from the network. At each iteration, one stimulation neuron received a two-pulse train with inter-pulse interval IPI_SNN_ = 3 ms (effective intra-train frequency 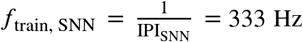). Pulses were modeled as brief suprathresh^I^o^PI^l^S^d^NN^current delta-like impulses applied to the selected neuron,

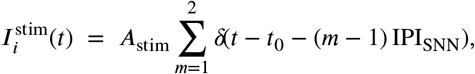

where *A*_stim_ is the supra threshold amplitude. After each two-pulse train, a recovery interval of *T*_inter-stim, SNN_ = 2 s was imposed before the next iteration. The procedure was repeated sequentially across all *M*_SNN_ stimulation sites and iterated for *N*_*T*_ = 200 trials per site. Spike trains from all *N*_SNN_ neurons were recorded continuously and analyzed with the same EC and IC pipelines used for the experimental data (Sections 4.3 and 4.3); unless otherwise noted, analysis windows and significance procedures matched the *in vitro* settings (see Supplementary Table 2 and Table S1b).

### 4.3 Causality Estimation

#### Interventional Connectivity

To quantify the causal impact of perturbing a source channel *i* on a target channel *j*, we compared the spiking activity of *j* before and after stimulation of *i* (Figure 1b, top central and right panels). Activity was evaluated within immediately pre- and post-stimulation time windows of equal duration *T*_*W*_ (*T*_*W*_ = 500 ms for experimental data, *T*_*W*_ = 300 ms for *in silico* simulations), chosen to balance statistical robustness and the (quasi) stationarity of neural dynamics within each window. In experimental data, to avoid contamination from electrical artifacts, short intervals surrounding the stimulation were excluded: the prestimulation window ended Δ_pre_ milliseconds before stimulation onset, and the post-stimulation window began Δ_post_ milliseconds after stimulation offset (see Figure S1a and Table S1b in Supplementary Information). For simulations, the corresponding intervals were negligible given the absence of hardware artifacts. Parameter values specific to each condition are reported in Supplementary Information.

For each stimulation site *i* and each recording channel *j*, we extracted all pre- and post-stimulation spike trains of *j* across the *N*_*T*_ repetitions of *i* stimulation, for time windows of length *T*_*W*_ (Figure 1b, bottom left and central panels). Within each window and trial, the spike train of *j* was divided into *n*_*w*_ bins (width *dt*_*w*_; see Table S1b), and the number of spikes in each bin was counted. This procedure yielded one set of spike counts for all pre-stimulation bins across all trials and another for all post-stimulation bins. Aggregating these values produced two empirical spike-count distributions for channel *j* conditioned on stimulation of *i*: (*p x*_*j*_| no stim) and (*p x*_*j*_| stim *i*). These distributions capture the variability of channel *j*’s spiking activity before and after perturbation of *i*. Their dissimilarity quantifies the causal impact of the stimulation. The interventional influence from channel *i* to channel *j* was then quantified as the statistical distance 𝒟 between these two distributions:

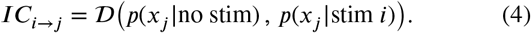

Among the available options for 𝒟, we adopted the Kolmogorov-Smirnov distance, which ranges from 0 (identical distributions) to 1 (maximally distinct distributions). A two-sided KS significance test (*α* = 0.05) was used to assess whether the difference between pre- and post-stimulation distributions was statistically significant under the null hypothesis of identical activity.

The resulting IC matrix (Figure 1b, bottom right panel) was constructed by evaluating *IC*_*i*→*j*_ for all source–target pairs, yielding an *M* × *N* array, where *M* is the number of stimulation sites and *N* the number of recording channels (or neurons in simulations). A summary of all parameter values used for experimental and simulated data is provided in Table S1b.

#### Effective Connectivity

We inferred the spontaneous causal flow among recorded channels by estimating the *Effective Connectivity* (EC) from ongoing spiking activity. Following Wiener’s (1956) definition of causality, EC quantifies how the past activity of a source channel *i* improves the prediction of a target channel *j*, beyond what can be explained by the target’s own past [32, 33]. Here, EC was not only used to describe spontaneous network organization, but also to assess its predictive power with respect to the evoked causal patterns measured through Interventional Connectivity (IC). In this study, EC was estimated using the *Transfer Entropy* (TE), a model-free, information-theoretic measure that captures directed dependencies between spike trains without assuming linearity in their interactions. For brevity, we use EC and TE interchangeably throughout the text, as TE provides the operational definition of EC in this work.

#### Data and preprocessing

TE was computed from spike trains recorded during spontaneous activity in the first experimental phase (Figure 1a). Spike times were discretized into bins of width *b* = 0.3 ms and binarized into time series *x*_*i*_(*t*) ∈ {0, 1} for each recording channel. This bin width was chosen to match the expected synaptic and axonal transmission times across the spatial scale of HD-MEA (a few millimeters), ensuring sensitivity to biologically plausible propagation delays [78]. In simulated data, no binning was applied, as spike times were already generated with a fixed simulation step Δ*t*_SNN_ = 1 ms; the resulting time series were likewise binarized to indicate spike occurrences at each timestep.

#### Transfer Entropy estimation

Delayed TE quantifies the directed information flow from a source channel *i* to a target channel *j* as:

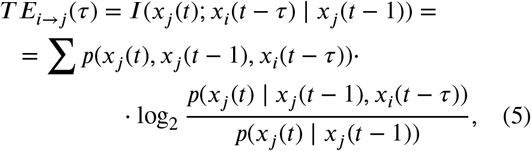

where *I*(*⋅*) denotes the conditional mutual information, and *τ* the time delay between source and target.

In experimental data, TE was computed for delays ranging from *b* to *τ*_max_ = 6 ms in increments of *b*, consistent with biologically plausible axonal propagation times. In *in silico* simulations, delays were explored in the range 1–10 ms to account for longer communication latencies in the model. For each source–target pair (*i, j*), the maximum TE value across delays,

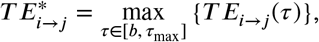

was taken as the effective connection strength, and the corresponding delay *τ*^∗^ as the estimated transmission latency (see Figure 1a).

#### Construction of the EC matrix

TE was computed from the binarized spike trains for all channel pairs (*i, j*), producing a full *N*×*N* matrix of directed effective connections (see Figure 1a right panel).For direct comparison with the experimentally derived IC matrix, the EC matrix was then reduced to an *M*×*N* subset, retaining only those links corresponding to stimulation–recording pairs probed during the IC protocol. The resulting EC matrix thus contained one directed weight 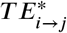 for each source–target pair, representing the maximum information transfer across delays. A summary of parameter values used for both experimental and simulated data is reported in Supplementary Information.

#### Significance test

To identify statistically meaningful interactions, we applied a surrogate-based significance test controlling for spurious coincidences, intrinsically caused by firing rates. For each source channel *i*, spike times were jittered by random displacements drawn from a Gaussian distribution (standard deviation 1 ms), while target spike times were left unchanged [34]. This preserved the burst and rate statistics of each channel, while disrupting directed dependencies between channel pairs. For each pair (*i, j*), TE was recomputed on a 1000-surrogate dataset; the connection was considered significant if the empirical value exceeded the 95^th^ percentile of the surrogate distribution (one-sided test, *α* = 0.05). Non-significant connections were set to zero.

### 4.4 Analytical Tools

#### Distance-matched null model for EC-IC correspondence

To test whether the correspondence between EC and IC could be explained by inter-electrode distance alone, we constructed a distance-matched null model. For each stimulation channel *i*, we quantified the observed reconstruction accuracy as the row-wise correlation *ρ* _*i*_ = corr (EC_*i*→∗_,IC_*i*→∗_) (Spearman correlation unless otherwise stated), computed across all recording channels *j* with finite values.

We then generated a null distribution that preserves the distance dependence of IC while destroying any EC-IC alignment beyond distance. Let *D*_*i*→*j*_ denote the Euclidean distance between stimulation site *i* and recording channel *j*. For each culture, we partitioned all off-diagonal distances into 15 distance bins using quantile edges, ensuring comparable occupancy across bins while maintaining sufficiently large within-bin target sets for within-row permutations. Within each stimulation row *i*, IC values were independently permuted among recording channels *j* falling in the same distance bin, leaving EC fixed. This procedure preserves, for each *i*, the empirical distribution of IC values at each distance scale, and therefore the IC spatial footprint expected from distance alone.

Repeating the distance-stratified shuffling for *N*_perm_ permutations yielded a null distribution 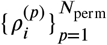 for each stimulation channel. We computed one-sided empirical *p*-values as 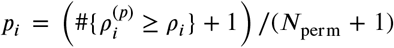, and controlled for multiple co^*i*^mparisons across stimulation channels using the Benjamini-Hochberg procedure at false discovery rate *α*.

#### Edge-level separability test (EC conditioned on IC significance)

For each stimulation source *i*, we tested whether outgoing EC weights were larger on IC-significant targets than on IC-non-significant targets. We split outgoing EC values into two sets using IC significance on the same edges (*α* = 0.05): IC-significant (responsive receivers) and IC-non-significant (non-responsive receivers).

We quantified separability with the difference between medians: 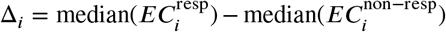. Significance was tested with a one-sided permutation test (*H*_0_ : Δ = 0, : Δ 0), preserving group sizes and using 20,000 permutations: 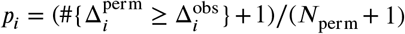. As a nonparametric effect size, we report Cliff’s delta and its probability-of-superiority transform, *P S*_*i*_ = (*δ*_*i*_ + 1)/2, which can be interpreted as the probability that a randomly chosen outgoing EC value from an IC-responsive target exceeds a randomly chosen outgoing EC value from an IC-non-responsive target. For each channel, we also computed a 95% bootstrap CI for Δ_*i*_ (5, 000 resamples; percentile CI, within-group resampling).

#### Shortest-path analysis

In simulated networks, shortest-path (SP) lengths were computed on the full *N* × *N* ground-truth connectivity matrix *W*_eff_, defined as the elementwise product between the structural weight *w*_*ij*_ and the synaptic gain *J*_*i*_. Edge distances were defined as *δ*_*ij*_ = 1/|*W*_eff,*ij*_| for all existing connections, such that stronger effective links corresponded to shorter distances. Two equivalent implementations were tested: one using the absolute values of *W*_eff_, and one restricted to the excitatory subnetwork obtained by setting negative weights to zero. Shortest paths were computed via Dijkstra’s algorithm, yielding a complete *N* × *N* matrix of pairwise path lengths. For direct comparison with the experimental IC and reduced EC matrices, the resulting SP matrix was downsampled to an *M* × *N* format, retaining only paths originating from the *M* stimulation sites to all *N* recording nodes.

## Acknowledgments

E.T. was supported from MUR and EU-FSE through the PhD fellowship PON Research and Innovation 2014-2020 (D.M 1061/2021) XXXVII Cycle / Action IV.4 “Tematiche Innovazione”. L.M. was partially supported by National Institutes of Health Awards R01NS118461, R01MH127375 and R01DA055439 and National Science Foundation CAREER Award 2238247. P.P. and S.V. were supported from NEUREKA project (EU H2020, FET, GA-863245). M.A. was supported by MUR, PRIN grant 2022HSKLK9, CUP C53D23000740006, within the project “Unveiling the role of low dimensional activity manifolds in biological and artificial neural networks”.

## Data and Code Availability

Effective connectivity was computed using the software package originally developed by [33] and adapted here for the purposes of this study: https://github.com/elisatentori/EC_calculation.

The SNN model simulation and the *in vitro* and *in silico* analysis scripts were performed using the package https://github.com/elisatentori/causality, released with this paper. Data are available upon reasonable request.

## CRediT authorship contribution statement

**Elisa Tentori:** Conceptualization, Methodology, Software, Visualization, Project administration, Writing – original draft. **George Kastellakis:** Methodology, Software, Writing - review and editing. **Marta Maschietto:** Methodology, Writing - review and editing. **Alessandro Leparulo:** Methodology, Writing - review and editing. **Panayiota Poirazi:** Funding acquisition, Methodology, Writing - review and editing. **Davide Bernardi:** Methodology, Writing - review and editing. **Luca Mazzucato:** Supervision, Conceptualization, Project administration, Writing - review and editing. **Michele Allegra:** Supervision, Conceptualization, Project administration, Writing - original draft. **Stefano Vassanelli:** Funding acquisition, Supervision, Conceptualization, Project administration, Writing - review and editing.

## Supplementary Information

### 1. Focal stimulation protocol and parameters

We used a standardized single-site stimulation protocol for all IC measurements, consisting of 200 trials per stimulation channel with fixed inter-stimulus intervals. The temporal structure of the protocol is shown in Figure S1a. The specific parameters employed *in vitro* and *in silico* are reported in Figure S1b. *In vitro* values were selected based on standard HD-MEA stimulation practices and were validated for robustness in the analyses reported in the next section, where we systematically varied *T*_*w*_, *dt*_*w*_, and Δ_post_ to assess the stability of IC reconstruction.

**Figure S1:**
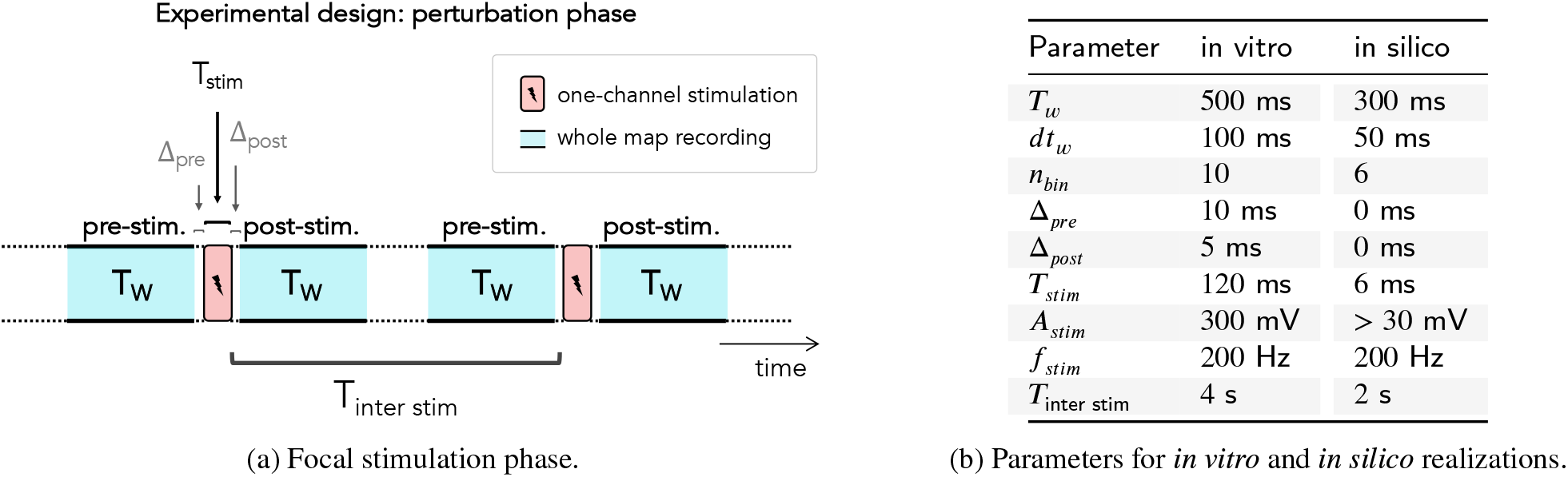
Focal stimulation phase: design and parameters.

### 2. Stability of spontaneous activity during EC estimation

Dissociated hippocampal cultures usually exhibit episodic network bursting, and slow fluctuations in overall firing rate are common [79]. We verified whether spontaneous activity used for EC estimation remained stationary throughout the 30-minute baseline recording and whether the statistical structure of bursting was stable. Raster plots (Figure S2b_1_–b_2_) show sustained bursting activity throughout the session. Culture 1 shows spatially clustered recruitment consistent with its modular organization (Figure S2a_1_), whereas Culture 2 exhibits spatially uniform activation. Spike-count distributions (Figure S2c_1_–c_2_) maintain the same heavy-tailed profile in the first and last 5 minutes, indicating preserved synchrony structure. Burst-duration distributions were statistically indistinguishable across the two intervals (Figure S2d_1_–d_2_; KS tests *p* = 0.19 and *p* = 0.65). Inter-burst interval (IBI) distributions likewise overlapped (Figure S2e_1_–e_2_), showing that the temporal organization of bursting did not change over the recording.

#### TE is stable as an edge-level estimate

We assessed the stability of TE as a function of recording duration and examined the structure of TE-edge distributions across cultures. In Culture 1, which exhibits modular organization, TE-edge distributions were bimodal (Figure S3a_1_, left): the higher mode corresponds to channel pairs within the same spatial module, whereas the lower mode reflects inter-module pairs. In contrast, the more spatially uniform Culture 2 displayed a unimodal distribution (Figure S3a_2_, left). In both cultures, TE values decreased systematically with inter-electrode distance (Figure S3a_1_-a_2_, right panels), consistent with spatially structured effective interactions. To evaluate sampling requirements, we quantified the convergence of TE estimates as a function of baseline recording duration. Spearman correlations between TE computed from progressively longer segments and the full 30-minute TE increased monotonically with sample size and saturated after approximately 10–15 minutes (Figure S3b), indicating that TE can be reliably estimated from limited data. Finally, we assessed EC–IC correspondence: spearman correlation remained stable across durations in Culture 1 (mean across channels: from *ρ*_5 min_ = 0.73 ± 0.12 to *ρ*_30 min_ = 0.75 ± 0.12) and increased with sample size in Culture 2 (mean across channels: from *ρ*_5 min_ = 0.42 ± 0.16 to *ρ*_30 min_ = 0.47 ± 0.13; Figure S3c). These results show that TE provides a robust, sample-efficient proxy for stimulation-evoked influence.

#### TE is robust as a source-level predictor

We then collapsed outgoing connectivity to a single summary value per stimulation channel. Comparable EC-IC correspondence was obtained when source output was summarized either by the median of outgoing links or by the mean of non-zero outgoing links, and the result was similar when EC was estimated from 15 or 30 min of spontaneous activity (Figure S4). Thus, the source-level ranking of influential stimulation sites is stable across reasonable choices of summary statistic and recording duration.

#### TE is informative enough to separate responsive from non-responsive targets

Finally, we asked whether EC and IC were aligned not only at the source level, but also in their edge-wise separation of responsive versus non-responsive targets. For each fixed stimulation channels in both cultures, recording channels classified as IC-responsive had systematically larger outgoing EC values than IC-non-responsive channels (see examples in Figure S5, top left). The converse analysis gave the same result: channels classified as EC-responsive showed systematically larger IC values than EC-non-responsive channels (Figure S5, bottom left). This effect is generalized across stimulation sites, as shown by the source-wise Probability of Superiority distributions, which were consistently shifted above chance in both cultures (Figure S5, right).

**Figure S2:**
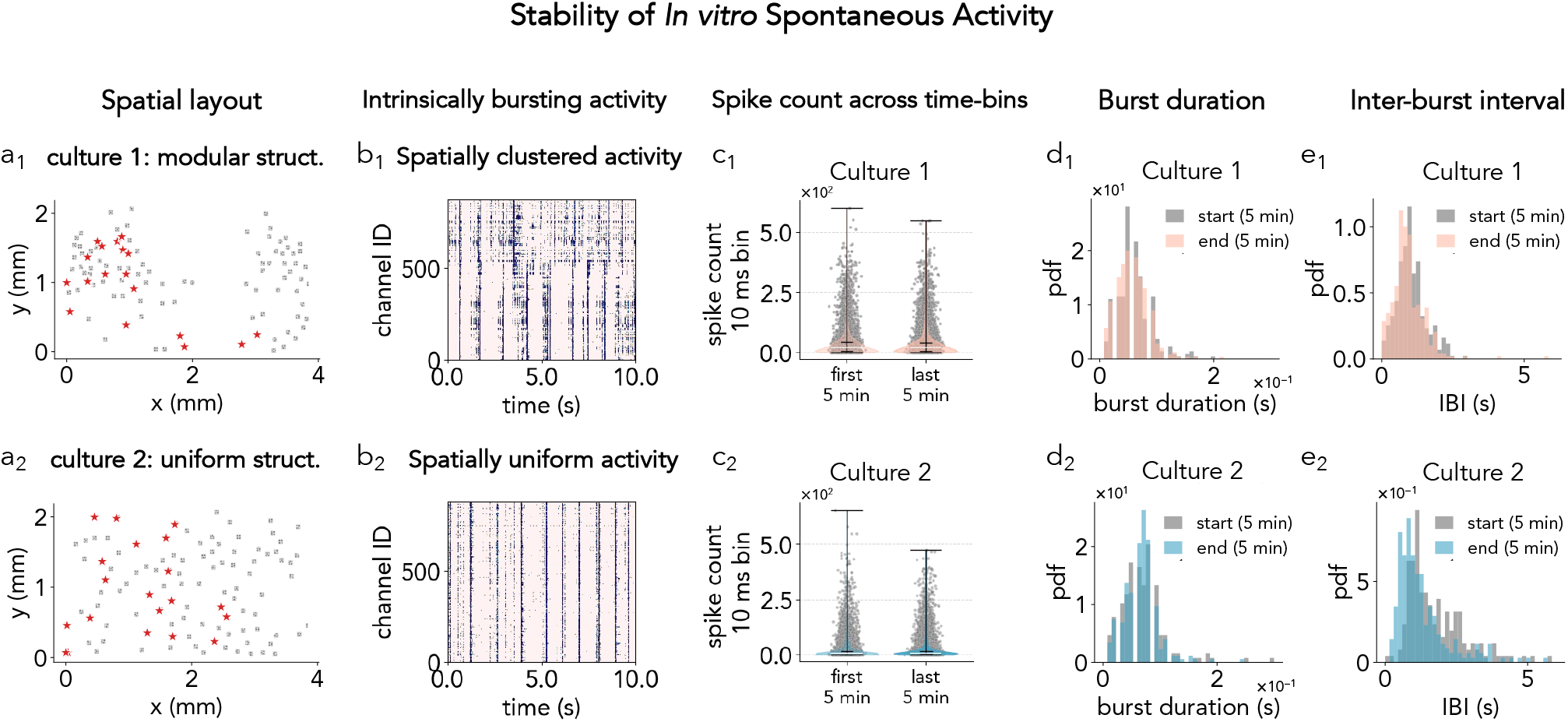
Stability of spontaneous activity during the EC experimental block. **(a**_1_,**a**_2_) Spatial layout of the *N*-channel recording map for two hippocampal cultures. Gray dots mark all recorded channels, red stars the *M* stimulation channels, and blue circles the example recording channels for which responses are shown in later panels. Culture 1 (a_1_) exhibits a modular spatial organization, whereas Culture 2 (a_2_) shows a more uniform distribution. **(b**_1_,**b**_2_) Raster plots showing persistent network bursting. Activity in Culture 1 (b_1_) depends on the channels position: intra-cluster activity is synchronized. **(c**_1_,**c**_2_) Spike-count distributions (10-ms bins) from the first and last 5 minutes show stable heavy-tailed profiles characteristic of synchronous bursting. **(d**_1_,**d**_2_) Burst-duration distributions in the first vs. last 5 minutes show no significant differences (Culture 1: KS = 0.087, *p* = 0.19; Culture 2: KS = 0.072, *p* = 0.65). Bursting dynamics remained stationary during the 30-minute EC recording. **(e**_1_,**e**_2_) Inter-burst interval distributions are likewise stable (KS < 0.10, *p* > 0.20 for both cultures).

**Figure S3:**
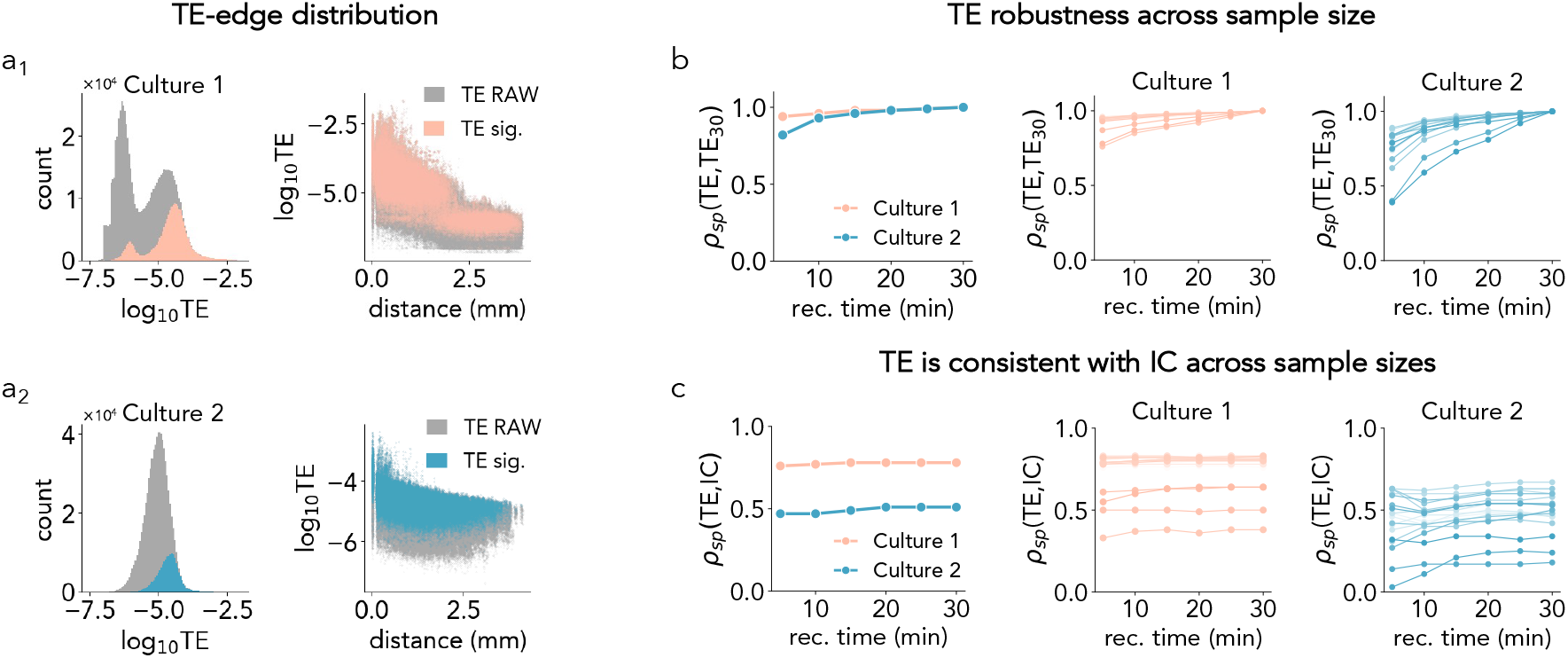
TE statistics, sampling robustness, and stability of EC–IC correspondence. **(a**_1_,**a**2) TE-edge distributions (log 10 scale) for Culture 1 and Culture 2. Gray indicates all TE values; colored distributions indicate statistically significant TE edges. Left, Culture 1 shows a bimodal distribution consistent with modular organization, whereas Culture 2 shows a unimodal distribution consistent with a more homogeneous spatial layout. Right, in both cultures, TE decreases with inter-electrode distance, indicating spatially structured effective interactions. **(b)** Convergence of TE estimates with recording duration. Spearman correlation between the TE matrix estimated from progressively longer recording segments and the reference TE matrix estimated from the full 30-min baseline increases monotonically and approaches saturation after ~ 10-15 min, indicating sample-efficient TE estimation. **(c)** Stability of EC-IC correspondence across recording durations. Source-wise Spearman correlation between TE and IC remains nearly constant across durations in Culture 1 and increases with sample size in Culture 2, showing that EC–IC correspondence is preserved even when TE is estimated from reduced spontaneous recordings.

**Figure S4:**
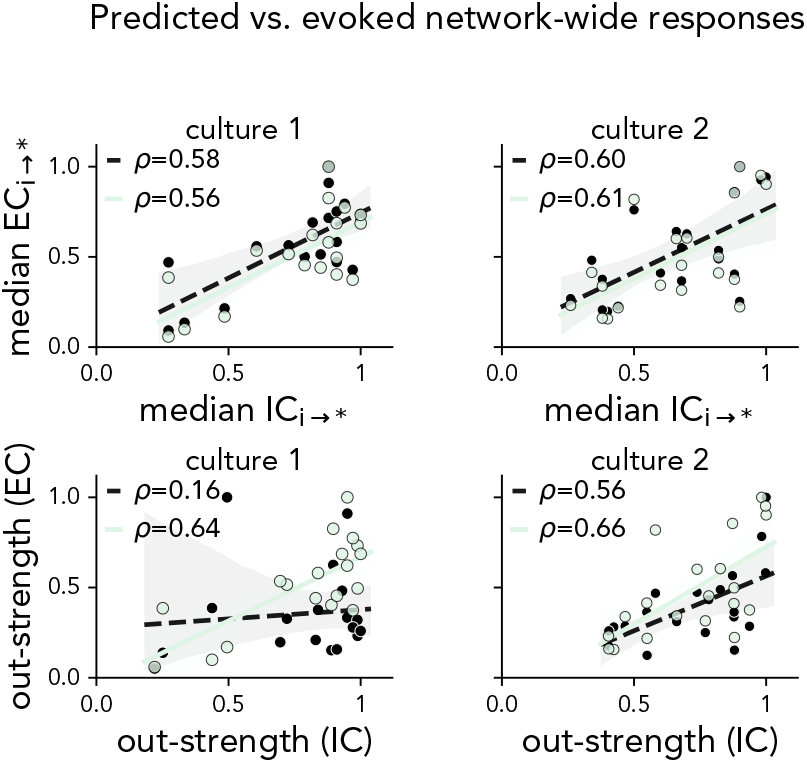
Source-level EC–IC correspondence is robust to recording duration and summary statistic. **Top:** correlation between median outgoing EC and median outgoing IC across stimulation channels, with EC estimated from 15 or 30 min of spontaneous activity. **Bottom:** same analysis when source output is summarized by the mean of non-zero outgoing links. Similar correlations across both recording durations and both summary definitions show that source-wise EC-IC correspondence is stable and does not depend on a specific choice of aggregation rule.

**Figure S5:**
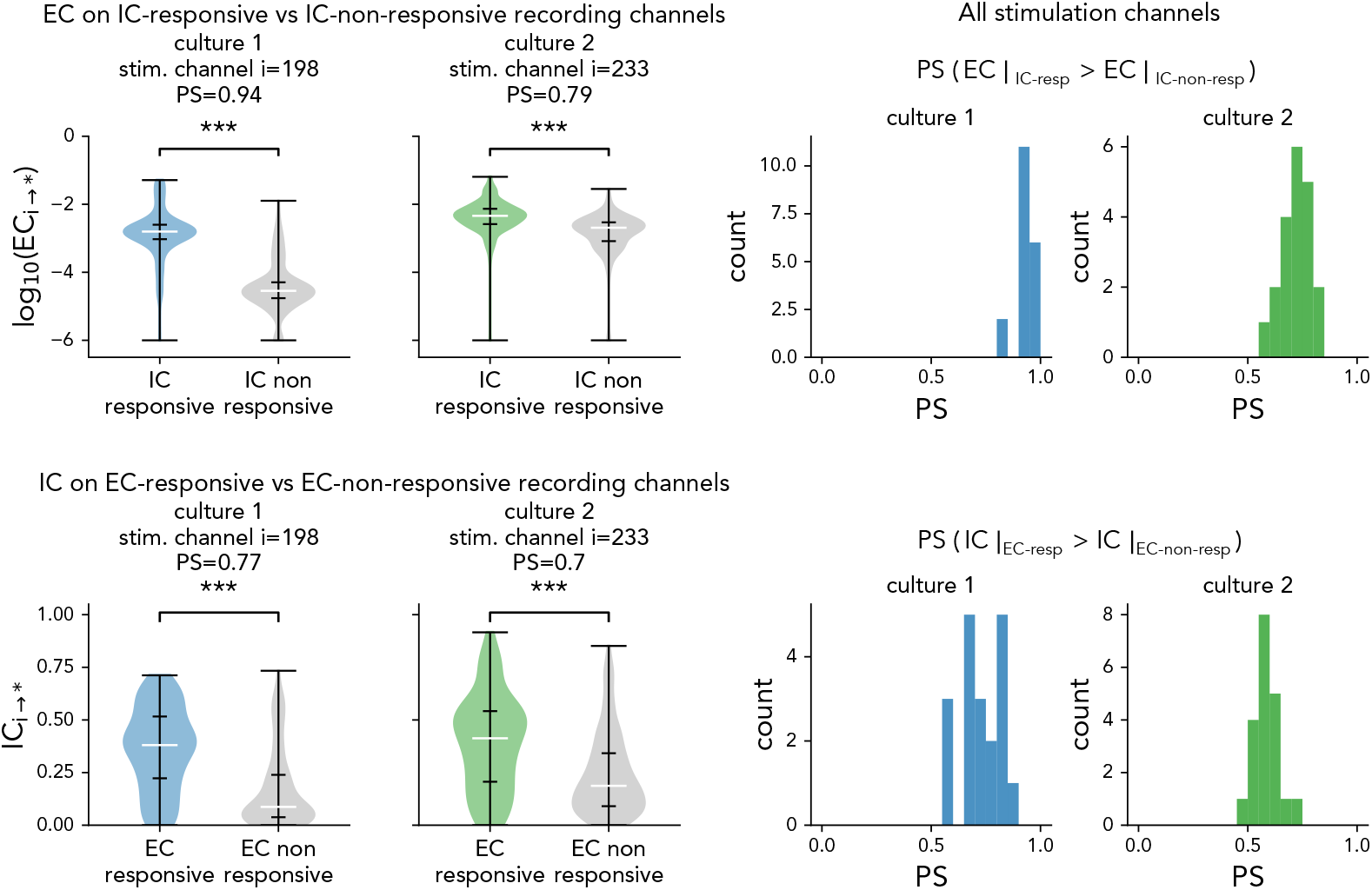
Edge-level separability analysis *in vitro*. **Top:** for one representative stimulation channel in each culture, outgoing EC values are compared between IC-responsive and IC-non-responsive recording channels; right, the corresponding source-wise Probability of Superiority (PS) across all stimulation channels. In both cultures, IC-responsive targets exhibit systematically larger EC values than IC-non-responsive targets. **Bottom:** inverse conditioning analysis. Outgoing IC values are compared between EC-responsive and EC-non-responsive recording channels for the same representative sources; right, the corresponding source-wise PS across all stimulation channels. EC-responsive targets likewise exhibit systematically larger IC values than EC-non-responsive targets. Together, the two analyses show reciprocal edge-level alignment between spontaneous EC and stimulation-evoked IC.

### 2. Robustness of IC estimation across window parameters

We quantified the robustness of IC with respect to the choice of *T*_*w*_, Δ_post_, and *dt*_*w*_, as well as to sample size (number of trials), and trial subsampling, and verified that the stimulation protocol did not induce detectable long-term plasticity (Figure S6).

First, we assessed TE–IC consistency across IC parameter choices. TE–IC correlations remained high across a broad range of pre/post window durations (*T*_*w*_) and post-stimulation delays (Δ_post_), and were similarly stable across bin sizes (Figure S6a, left and middle). TE–IC correspondence increased with the number of stimulation trials and saturated after ~ 100λ150 trials in both cultures (Figure S6a, right), confirming that TE provides a robust predictor of stimulation-evoked influence. Second, we quantified the internal robustness of IC itself. IC matrices computed with different window parameters were strongly correlated with the reference IC (*T*_*w*_ = 500 ms, Δ_post_ = 5 ms), with saturation reached for windows above ~100–150 ms (Figure S6b, left). IC estimates also became progressively more stable as the number of trials increased (Figure S6b, right), demonstrating that IC reconstruction is reliable and not sensitive to modest parameter variations. Third, we asked whether the TE–IC correspondence was stable across independent subsets of stimulation trials. For each culture, we recomputed IC using two disjoint halves of the trials and, for every stimulation site, quantified the TE–IC correlation on a row-by-row basis (i.e. by correlating the outgoing TE vector of each source channel with the corresponding IC vector). Split-half validation yielded nearly identical distributions of TE-IC correlations (Figure S6c), demonstrating that the alignment between TE and IC reflects reproducible response structure across trials rather than dependence on specific repetitions of the perturbation protocol. Finally, we verified that stimulation did not induce long-term reorganization of network activity. For each stimulation site, we compared post-stimulus activity vectors from the first and last 20 trials. Correlation matrices from early and late trials were nearly identical (Figure S6d, left and middle), and the correlation between early–late responses across all sites was high (*ρ*_pearson_ = 0.90; Figure S6d, right). These results rule out stimulation-induced plasticity over the timescale of the perturbation block and confirm that IC reflects stationary network dynamics.

### 4. Spiking Neural Network Model

To mechanistically validate the causal–inference framework, we implemented a minimal spiking network model that reproduces (i) the spontaneous bursting regime of dissociated hippocampal cultures and (ii) the stereotyped motifs evoked by focal electrical stimulation. The model integrates four essential components: (1) spatially embedded excitatory–inhibitory (E–I) connectivity, (2) Izhikevich single-neuron dynamics, (3) distance-dependent axonal delays, and (4) presynaptic shortterm synaptic depression (STD). As shown in the main text, this combination is sufficient to generate measurable poststimulation reverberation, and STD is necessary for the emergence of IC, despite unchanged structural connectivity and stimulation protocol.

#### Network Topology

We simulated a two-dimensional, spatially embedded E–I network of *N* = 1000 neurons with E/I ratio 4:1. Excitatory neurons comprised Regular Spiking (RS) and Intrinsically Bursting (IB) classes; inhibitory neurons comprised Fast Spiking (FS) and Low-Threshold Spiking (LTS). Neurons were arranged in a square domain (mm units) using Poisson–disk sampling, yielding near-uniform density and well-defined inter-neuronal distances *d*_*ij*_.

Connectivity was directed and sign-constrained (*E* → {*E, I*} > 0, *I* → {*E, I*} < 0). Excitatory connectivity followed an EDR:

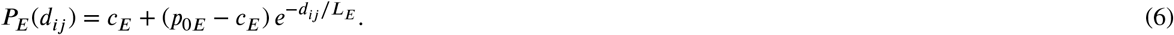

applied to both E and I targets; this yields dense local excitation with a controlled distal tail set by *L*_*E*_. Inhibitory projections were local:

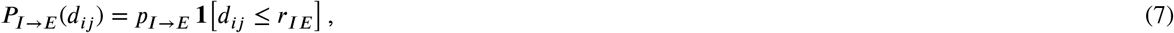

and if no excitatory target was available within *r*_*IE*_, the nearest excitatory neuron was connected.

Synaptic weights were drawn from Gaussian distributions with distance-dependent mean:

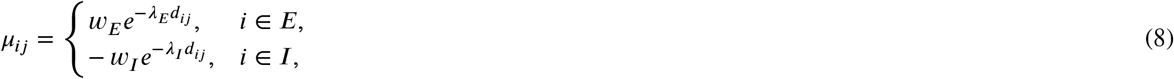

and standard deviation *σ*_*ij*_ = CV|*μ*_*ij*_|.

Communication delays were discretized in Δ*D* = 1 ms steps. Excitatory delays scaled with distance and ranged from 1–10 ms; inhibitory delays were fixed at 1 ms.

**Figure S6:**
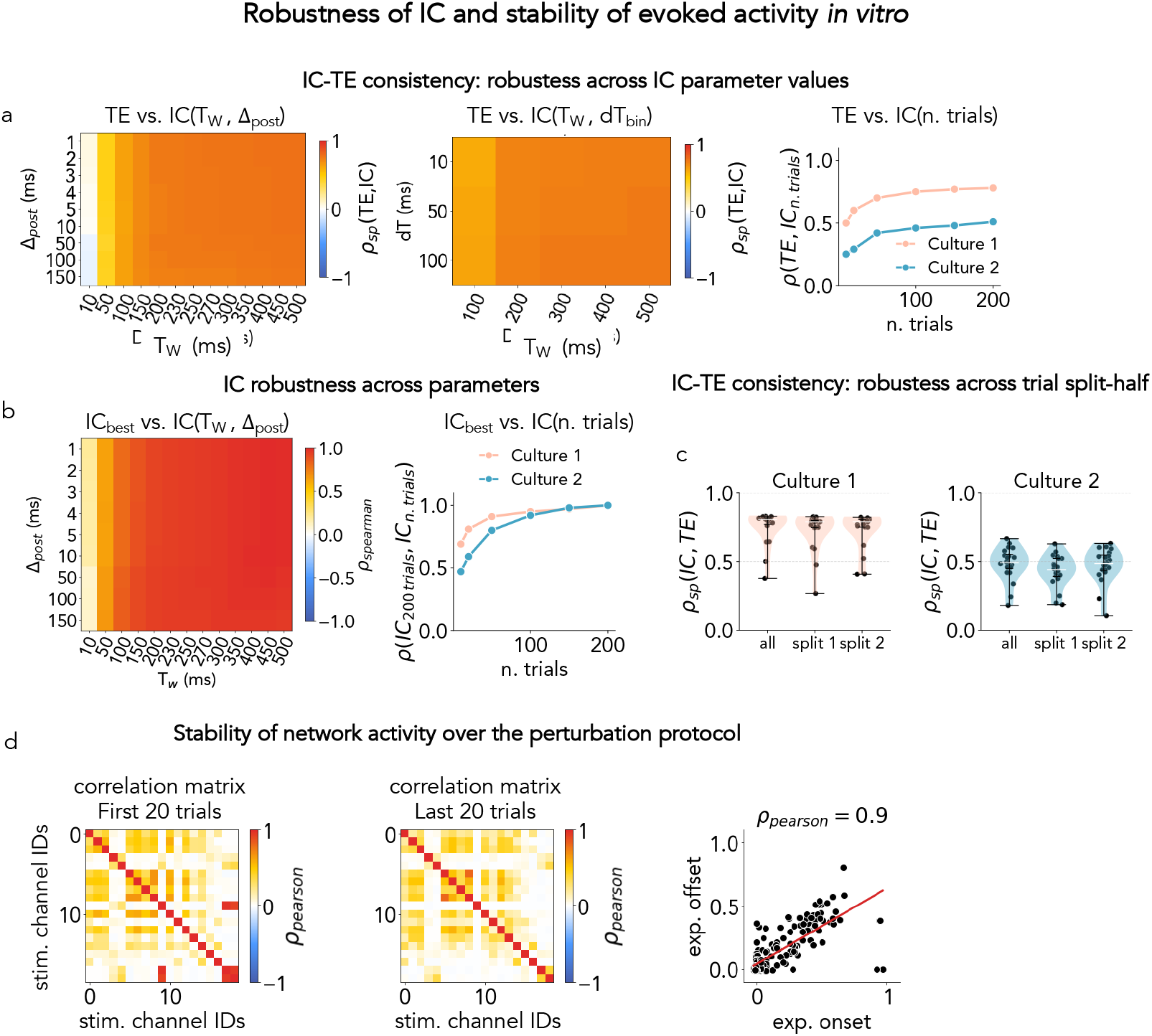
Robustness of IC estimation and stability of evoked activity *in vitro*. **(a)** TE–IC correspondence across IC parameter choices and trial counts. **Left:** Spearman correlations between TE and IC edges across a grid of pre/post window parameters (*T*_*w*_, Δ_post_). **Middle:** Same analysis across different bin sizes (*dt*_*w*_). **Right:** TE–IC edge correspondence as a function of the number of stimulation trials, showing saturation for ~ 100 − 150 trials in both cultures. **(b)** IC robustness across window parameters and sample size. **Left:** Correlations between the reference IC (IC_*best*_) and IC computed with alternative (*T*_*w*_, Δ_post_) choices; reconstruction stabilizes for *T*_*w*_ ≳ 150 ms. **Right:** Correlation between IC computed from the first *n* trials and the full IC matrix, showing monotonic stabilization with trial count. **(c)** Split-half validation of TE–IC correspondence: for each stimulation site, TE–IC correlations were computed row-by-row using IC estimated from two disjoint sets of 100 trials. The two distributions closely match the full dataset for both cultures, indicating that TE–IC alignment is reproducible across independent subsets of trials. **(d)** Stability of evoked responses across the stimulation protocol. Correlation matrices of post-stimulation activity from the first 20 trials (left) and last 20 trials (middle) exhibit highly similar structure. **Right:** Early vs. late correlations across stimulation sites (*ρ*_pearson_ = 0.90), indicating no detectable long-term plasticity during the perturbation phase.

#### Network Dynamics

Each neuron follows the quadratic integrate–and–fire model with adaptation [71] and synaptic depression [35–38]. Single-neuron dynamics are described by the coupled differential equations:

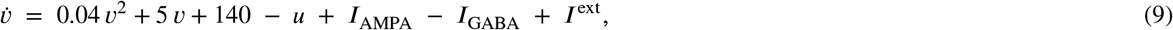

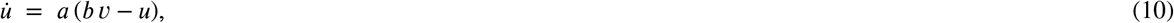

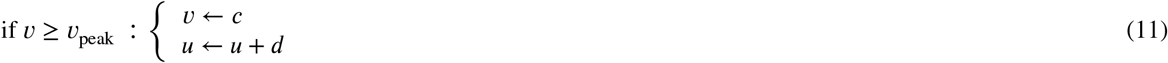

where *v* and *u* describe the membrane potential and the membrane recovery. When the membrane potential reaches *v*_peak_ = 30 mV, *v*(*t*) is reset to *c*, and *u*(*t*) is incremented by *d*, capturing the essential dynamics of action potential generation and recovery. Class–specific parameters *a, b, c, d* are used for each neuron type (RS, IB, FS and LTS; see [71, 72]), simulating the diversity of neuronal firing behaviors observed in biological systems. After each spike an absolute refractory period is enforced (excitatory 3 ms, inhibitory 1 ms), during which *v* is clamped to *c*.

Excitatory and inhibitory inputs are carried by AMPA and GABA currents that relax exponentially and receive delayed presynaptic impulses [37, 38, 73]. Let 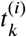 denote the *k*-th spike time of a presynaptic neuron *i*, and *D*_*ij*_ the axonal delay from *i* to a postsynaptic neuron *j* (integer mu^*k*^ltiples of Δ*D*). The postsynaptic currents at *j* satisfy

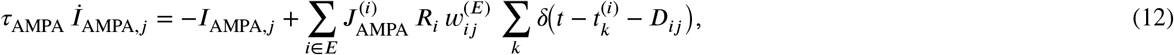

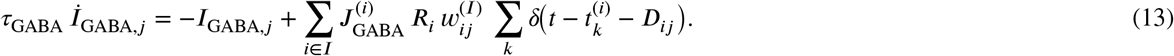

Here 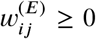 and 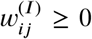 are non-negative synaptic strengths from *i* to *j* on AMPA and GABA channels; the net sign enters Eq. (9) via +*I*_AMPA,*j*_ and − *I*_GABA,*j*_. Presynaptic gains 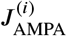 and 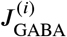 are drawn independently to capture release heterogeneity,

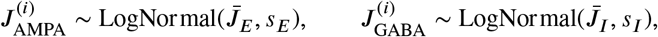

with target means 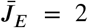 (excitatory → AMPA) and 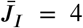 (inhibitory → GABA) (see Table 2 for parameters and Figure **??**F). These gains multiply the resource *R*_*i*_ and the synaptic strength *w*_*ij*_ in Eqs. (12)–(13), setting the per-spike impulse amplitude *J* ^(*i*)^*R*_*i*_*w*_*ij*_.

**Table 1.**
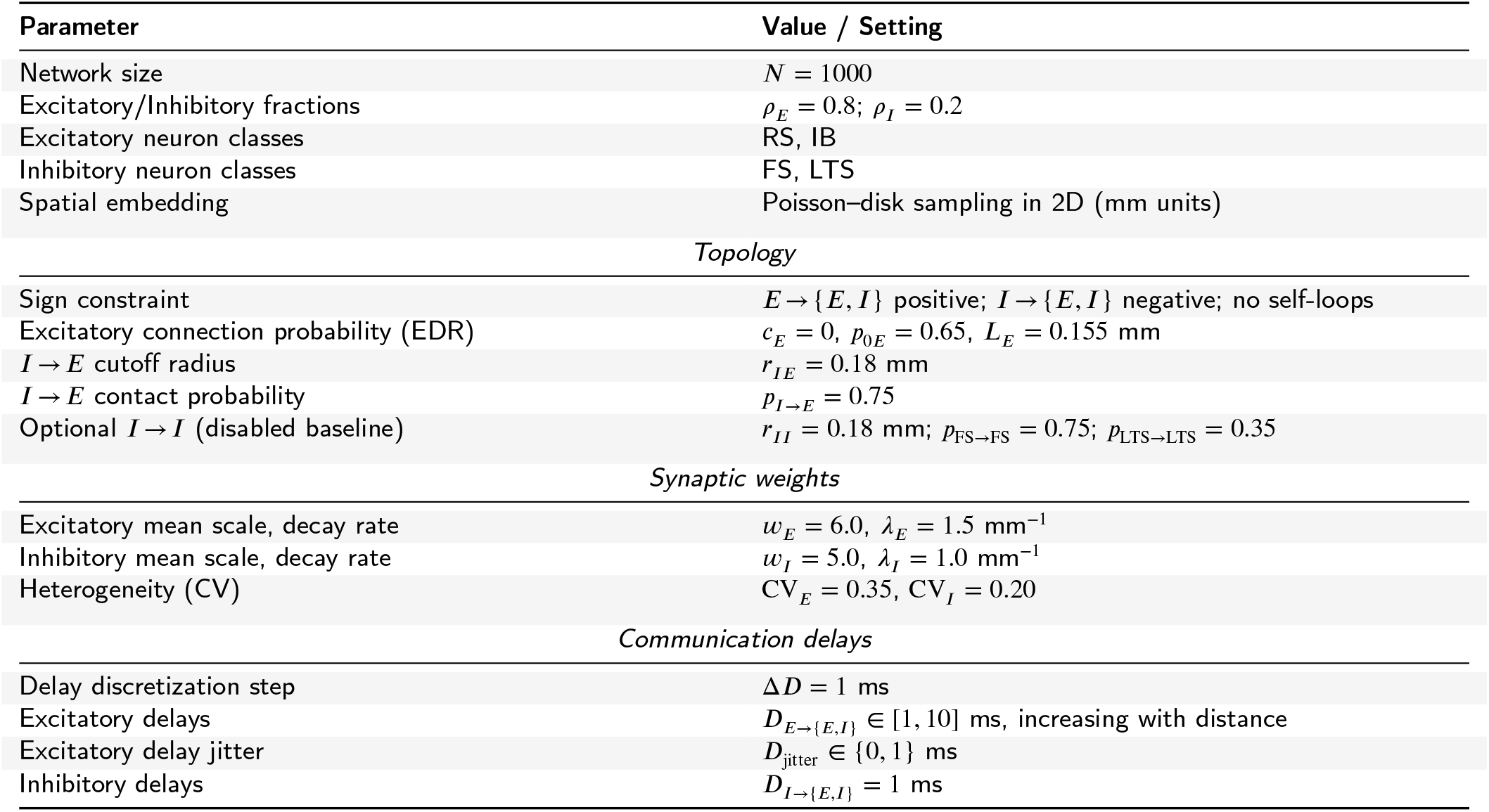
Parameters used to construct the spatially embedded excitatory–inhibitory network.

**Table 2.** Parameters for the spiking network with Izhikevich dynamics [71], Tsodyks–Markram depressing release [35, 36], excitatory Poisson shot noise, and delayed AMPA/GABA currents [37, 38].

| Quantity | Definition | Baseline value / note |
| --- | --- | --- |
| Spike peak | $v_{\text{peak}}$ | 30 mV |
| Izhikevich params | $a, b, c, d$ | class-specific (RS/IB for E; FS/LTS for I). See parameters in [71] |
| Simulation time step | $\Delta t$ | 1 ms |
| Absolute refractory | $\tau_{\text{ref}}^E, \tau_{\text{ref}}^I$ | 3 ms (E), 1 ms (I) |
| AMPA decay | $\tau_{\text{AMPA}}$ | 3 ms |
| GABA decay | $\tau_{\text{GABA}}$ | 50 ms |
| Resource recovery | $\tau_R$ | 500 ms |
| Depression factors | $\beta_E, \beta_I$ | 0.1, 0.8 |
| Presynaptic gains (E→AMPA) | $J_{\text{AMPA}}^{(i)}$ | lognormal, mean $\bar{J}_E=2$, $s_E=0.25$ (per presynaptic E) |
| Presynaptic gains (I→GABA) | $J_{\text{GABA}}^{(i)}$ | lognormal, mean $\bar{J}_I=4$, $s_I=0.25$ (per presynaptic I) |
| Synaptic strengths | $\tilde{w}_{ij}^{(E)}, \tilde{w}_{ij}^{(I)}$ | non-negative; sign via (9) |
| Delays | $D_{ij}$ | E: 1–10 ms (increasing with distance); I: 1 ms |
| Excitatory shot noise | $A_E, \nu_E$ | 13.5 mV, 1.2 Hz (applied to all neurons) |

Presynaptic release is modulated by a resource variable *R*_*i*_ ∈ [0, 1] per presynaptic neuron *i*, following the depression-only limit [37] of the Tsodyks–Markram model [36]:

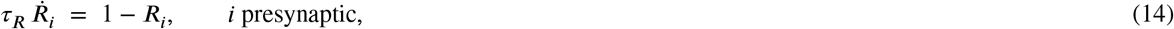

and at each presynaptic spike time 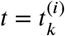 set

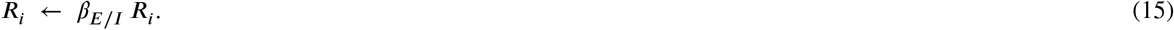

The depression time constant *τ*_*R*_ 0.5 second, making it the slowest timescale in the pulse dynamics [74]. In this parameterization, a presynaptic spike from *i* to *j* contributes an impulse of amplitude 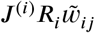 in (12)–(13); in Tsodyks–Markram notation [36] this corresponds to a fixed utilization *u* = 1 − *β*_*E*/*I*_ absorbed into *J*, leaving *R*_*i*_ as the sole dynamical resource governing short-term depression.

All neurons (E and I) receive an excitatory shot–noise drive,

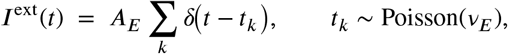

with population–wide rate *ν*_*E*_ and amplitude *A*_*E*_. This term represents effectively several elements that are not explicitly modeled, such as unresolved excitatory afferents from outside the simulated subnetwork as well as intrinsically active neurons that are needed to keep the network active [76, 77]. Background fluctuations play a role in shaping the network response to stimuli, and play a role in the amplification-stability tradeoff that enable downstream propagation of localized perturbations in cortical neuronal networks [80–82].

#### Control model without distance-dependent excitatory connectivity (no EDR)

As a control, we removed the EDR while keeping neuronal positions fixed (Figure. S7). In this condition, E, S and B motif-like single-target responses were still observed, but the dependence of motifs and of the perturbation effect (IC) on source–target distance largely disappeared in both topologies. Thus, EDR is not required for response motifs per se, but it is necessary to reproduce their spatial organization.

Consistently, removing EDR flattened the global spatial footprint of structure, EC, and IC (Figure S8a). Nevertheless, source-wise EC–IC correspondence remained above the distance-stratified null model, indicating that source-specific predictability can persist even when the distance-dependent excitatory connectivity is disrupted (Figure S8b).

#### Effect of presynaptic STD on post-stimulation reverberation

To test whether presynaptic STD can account for the emergence of post-stimulation reverberation in the model, we compared two otherwise identical networks: (i) a full model including presynaptic STD on all synapses, and (ii) a version in which STD was removed while keeping topology, neuron types, synaptic weights, delays, and external drive unchanged (Figure S9). In the model without STD, focal stimulation elicited only a brief, spatially confined response, and the corresponding IC matrix contained almost no detectable outgoing influence. In contrast, the model with STD produced extended post-stimulation activity and a structured IC, with propagation patterns spanning multiple network modules. Distance–dependent IC profiles confirmed this difference: without STD, IC decayed sharply with distance; with STD, the decay was broader and multi-peaked, consistent with recurrent propagation through the network. These results do not imply that STD is the biological mechanism underlying reverberation in dissociated cultures. Instead, we show that STD is a minimal and sufficient component within this modeling framework to reproduce the key phenomenology observed experimentally: reproducible post-stimulation reverberation, spatially structured IC, and alignment between IC and the underlying connectivity.

### 2. Alternative EC metrics

To assess the robustness of our conclusions beyond TE, we recomputed EC using two linear metrics that capture delayed spike-time dependencies: Signed Cross-Correlation (SC) and Cross-Covariance (XCov). These metrics were selected because they isolate rate-driven, time-lagged interactions under linear assumptions, thus providing complementary validation to the model-free TE framework.

**Figure S7:**
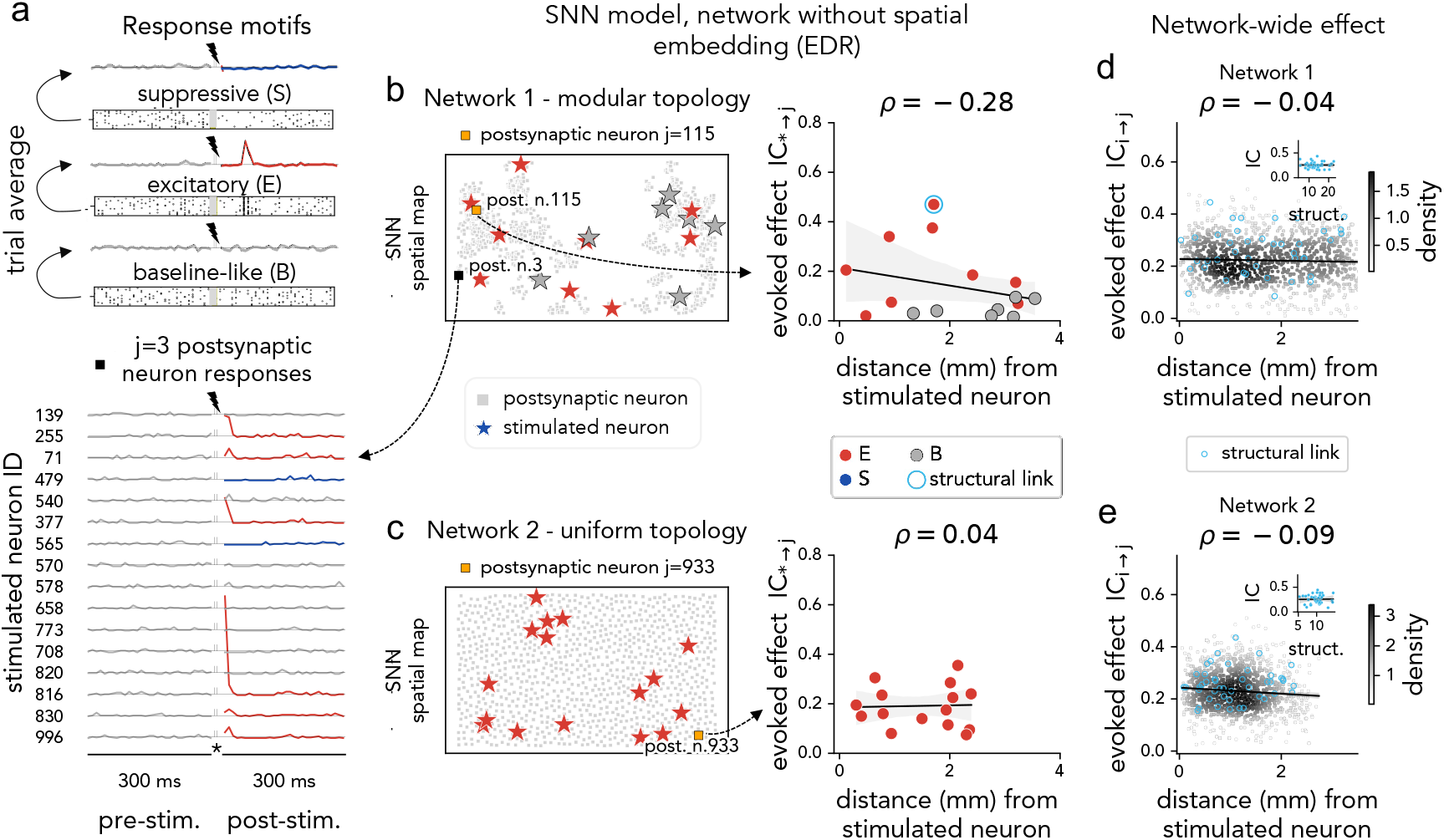
Control model without distance-dependent excitatory connectivity (no EDR): response motifs persist but lose their distance-dependent organization. **(a)** Definition of the three response motifs (S, E, B) and trial-averaged responses of one example postsynaptic neuron to stimulation from multiple source neurons. **(b**,**c)** Example source-specific responses in the modular and uniform networks. *Left:* spatial layout with stimulated neurons (stars; colored by recording channel response motif) and the selected postsynaptic neuron (orange square). *Right:* evoked effect *IC*_*i*→*j*_ versus Euclidean distance from the stimulated neuron. In the absence of EDR, the correlation between effect magnitude and distance is strongly reduced or absent. **(d**,**e)** Population-level IC across all stimulation–recording pairs for the two network topologies. The network-wide effect no longer shows the structured distance-dependent decay observed in the baseline model.

**Figure S8:**
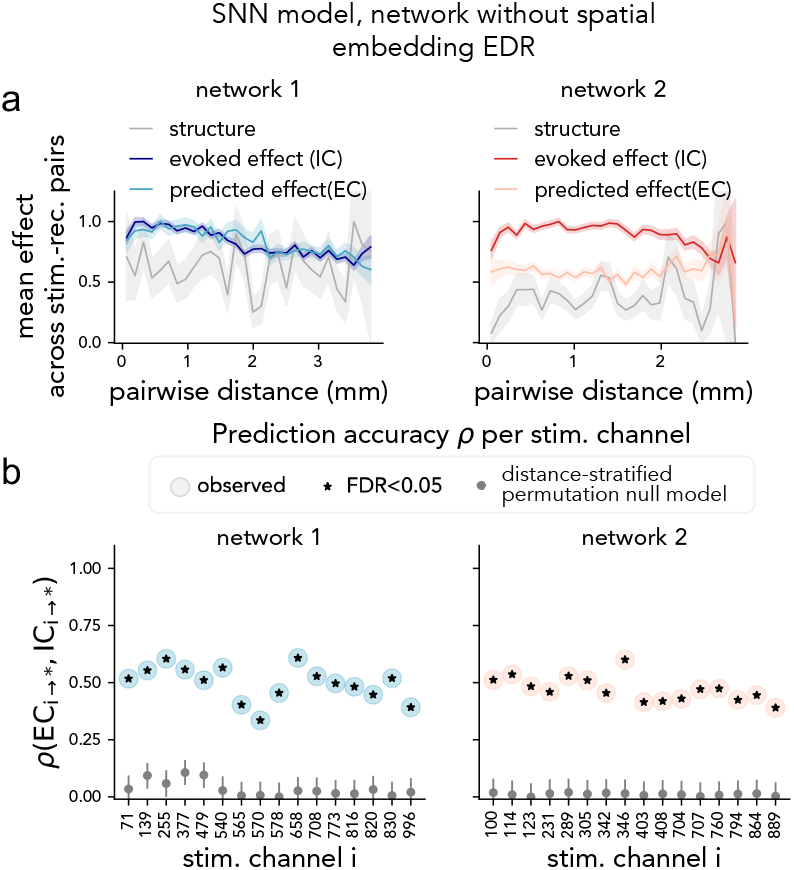
Control model without distance-dependent excitatory connectivity (no EDR): global spatial footprint and source-wise prediction. **(a)** Mean structural weight, mean evoked effect (IC), and mean predicted effect (EC) are plotted against pairwise distance for the modular (left) and uniform (right) networks. Compared with the baseline model, the spatial ordering of structure, EC, and IC is markedly weakened in the absence of EDR. **(b)** For each stimulated neuron, source-wise prediction accuracy *ρ*(*EC*_*i*→∗_, *IC*_*i*→∗_) is compared with a distance-stratified permutation null model. Although the spatial footprint is disrupted, EC–IC correlations remain above the null distribution, indicating residual source-specific correspondence not explained by distance alone.

**Figure S9:**
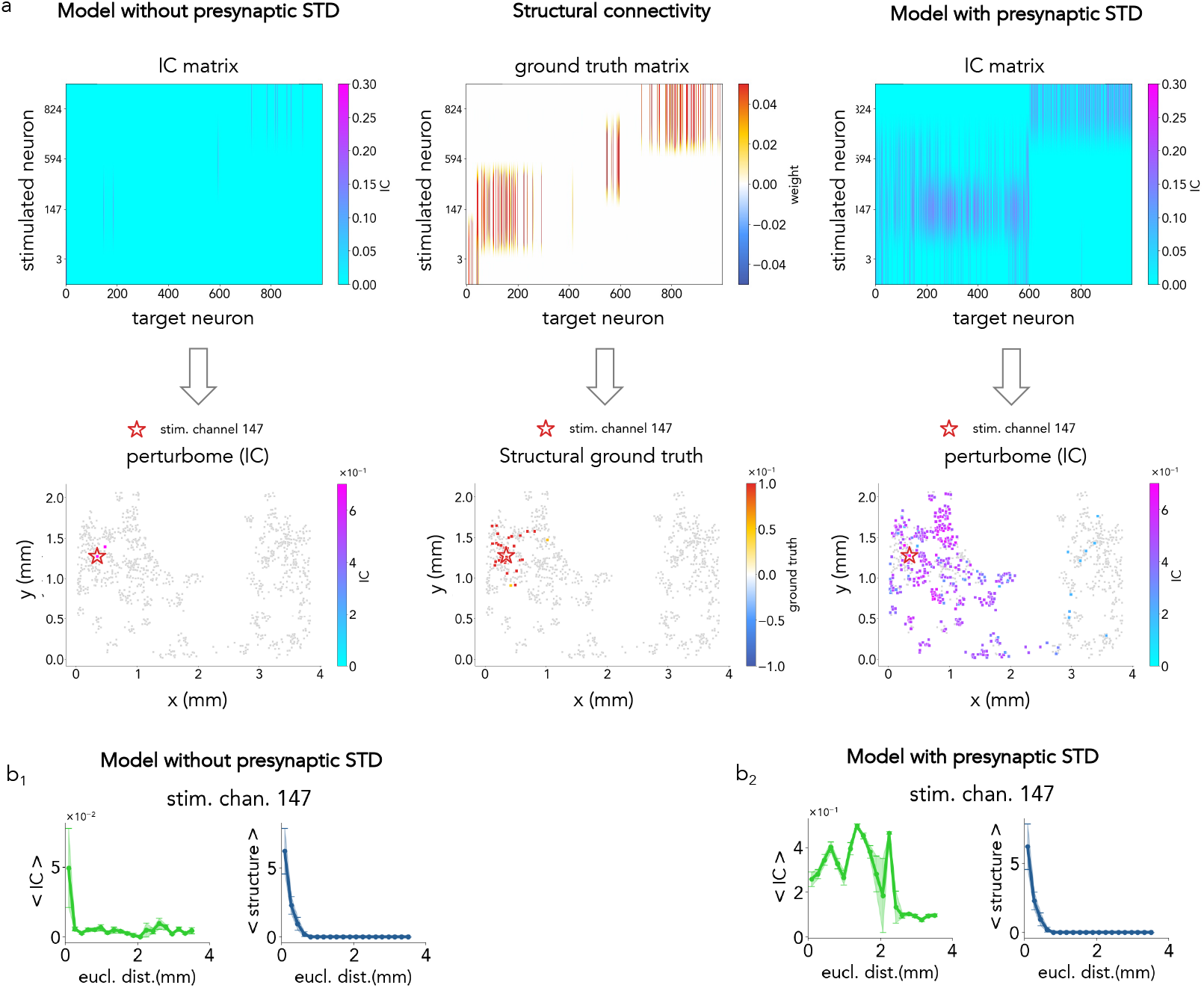
Effect of short-term synaptic depression (STD) on post-stimulation reverberation in the model. **(a)** IC matrices for the model without STD (left), the structural connectivity (middle), and the model with STD (right). Removing STD suppresses reverberation: IC remains sparse and confined to the stimulated site. Including STD yields structured, spatially extended IC that reflects the underlying connectivity. **(b)** Distance–dependent IC profiles for the same stimulation site. **b**_**1**_ Without STD, IC decays sharply with distance. **b**_**2**_ With STD, IC displays a broader profile with secondary peaks, reflecting propagation through recurrent pathways. These results indicate that STD is sufficient, within this model, to generate reverberation and reproduce the spatial IC structure seen *in vitro*.

All EC estimates were computed from *T*_*rec*_ = 30 minutes of spontaneous activity (bin size *b* = 0.3 ms), following exactly the same pipeline used for TE: delayed coincidence curves were evaluated over *τ* ∈ [0, 5] ms, significance was assessed via identical spike-time jittering, and a firing-rate–independent normalization was obtained by *Z*-scoring relative to the jittered null distributions. The resulting EC matrices were then restricted to the *M* × *N* subset corresponding to experimentally measured IC.

#### Signed Cross-Correlation

Signed Cross-Correlation (SC) [83] measures deviation of the cross-correlogram from its mean across delays, thereby removing coincidence baselines due to firing rates. Positive peaks reflect excitatory interactions and negative peaks inhibitory ones, with connection strength given by the maximum absolute SC value.

#### Cross-Covariance

Cross-Covariance (XCov) quantifies the covariance between *x*_*t*_ and *y*_*t*−*τ*_ after subtracting the expected coincidence count under independence. The peak XCov value across delays was used as the connection estimate.

#### Rationale on the metrics

Although TE captures nonlinear interactions while SC and XCov reflect linear dependencies with distinct baseline corrections, all three metrics produce comparable large-scale EC structure. This supports the robustness of the spontaneous-to-evoked correspondence described in the main text.

#### Alternative EC Metrics: Results

To assess whether the EC–IC correspondence depends on the specific choice of EC metric, we repeated the full analysis using also SC and XCov. All three metrics exhibited comparable spatial footprints, with clear distance-dependent decay (Figure S10c), and all showed a similar monotonic relationship with IC (Figure S10b). For each stimulation site, we quantified the alignment between spontaneous and evoked influence by computing the Spearman correlation between its outgoing EC and IC profiles (Figure S10d_1_,d_2_). Across cultures, TE, SC, and XCov produced nearly indistinguishable EC–IC similarity maps. These results indicate that the predictive relationship between spontaneous activity and perturbation-evoked influence is not specific to TE. Despite relying on different assumptions (nonlinear vs. linear), TE, SC, and XCov provide consistent estimates of directed interactions relevant to IC. This robustness supports the interpretation that the EC–IC correspondence reflects a genuine circuit-level organization rather than a metric-specific artifact.

**Figure S10:**
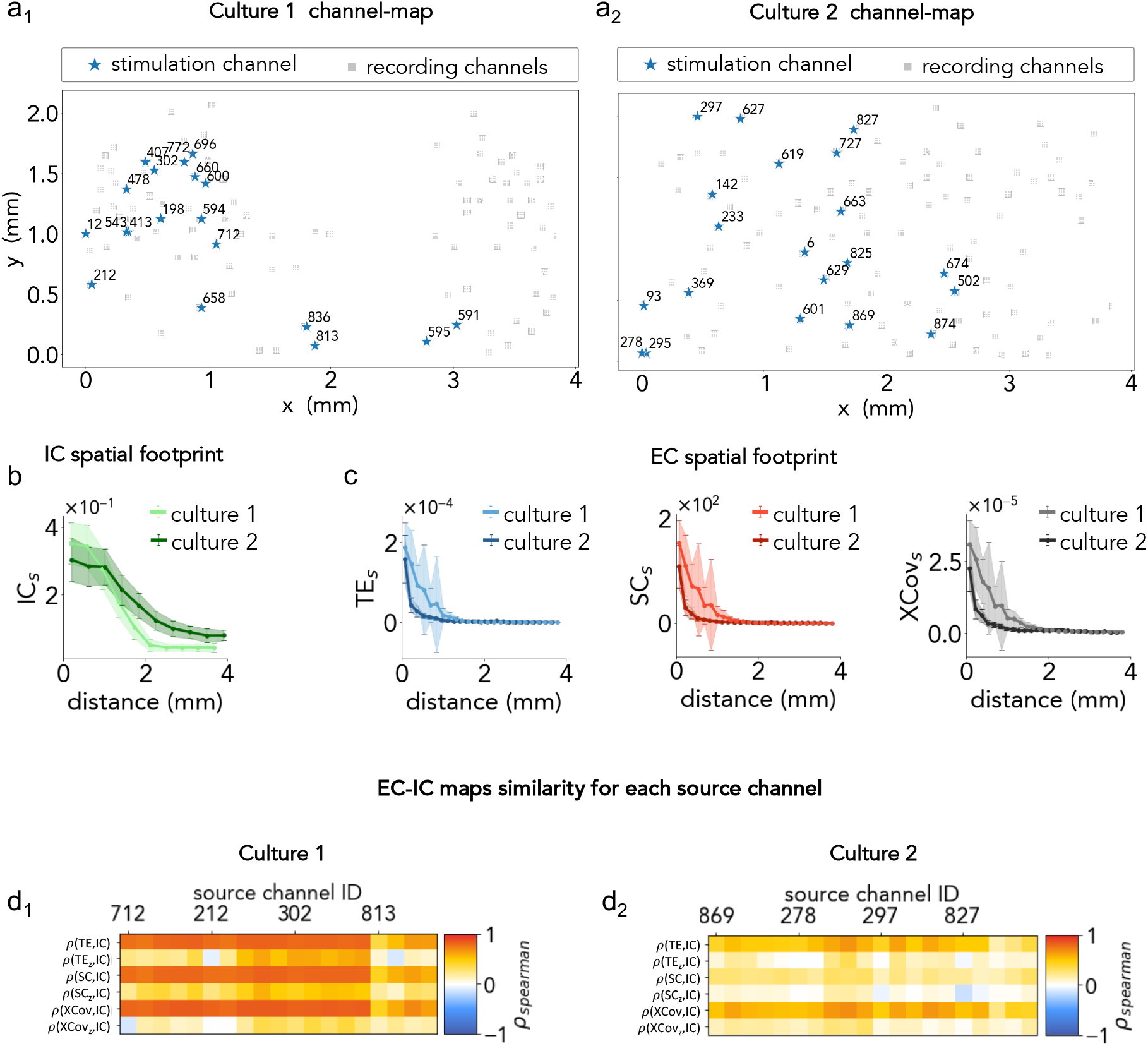
Comparison between IC and multiple EC metrics computed from spontaneous activity. **(a**_1_**–a**_2_) Spatial layout of stimulation (blue star) and recording channels (gray squares) for Culture 1 and Culture 2. **(b)** Spatial footprint of IC, showing distance–dependent decay of the evoked response. **(c)** Spatial footprints of the three EC metrics (TE, SC, XCov) for both cultures. **(d**_1_**–d**_2_) EC–IC similarity for each source channel. Each column corresponds to a stimulation site; each row reports the Spearman correlation between IC and one EC metric (TE, SC, XCov and z-scored versions TE_*z*_, SC_*z*_, XCov_*z*_) computed on its outgoing links.

### 7. Signed Interventional Connectivity

The standard IC measure captures the magnitude of stimulation-evoked influence but not its sign. To test whether the direction of the response could be recovered, we constructed a signed version of IC.

We shortened the pre- and post-stimulation windows to *T*_*w*_ = 50 ms to isolate the early component of the response. For each stimulation–recording pair, we computed two one-sided KS statistics: one testing whether the post-stimulation spike-count distribution exceeded the pre-stimulation one, and one testing the opposite. The signed IC value was defined as the statistic with the largest absolute magnitude, with the sign indicating the direction of the change. The resulting matrix (Figure S11b) closely matched the raw post–pre spike-count differences (Figure S11a), and the one-sided KS behaved as the signed analogue of the standard two-sided KS (Figure S11c).

We then asked whether signed EC measures computed from spontaneous activity (specifically SC and XCov without taking the absolute value) could recover the sign of IC. Both metrics showed weak or negative correlations with signed IC (Figure S11d–e, left). In contrast, their unsigned variants retained moderate correspondence with the unsigned IC magnitude (Figure S11d–e, center and right). Thus, spontaneous activity provided reliable information about the *strength* of directed interactions but not about their *sign*. The early stimulation-evoked response often contained inhibitory components (probably the activation of recurrent inhibitory circuits) that were not detectable in spontaneous dynamics, preventing signed EC metrics from predicting the direction of IC.

**Figure S11:**
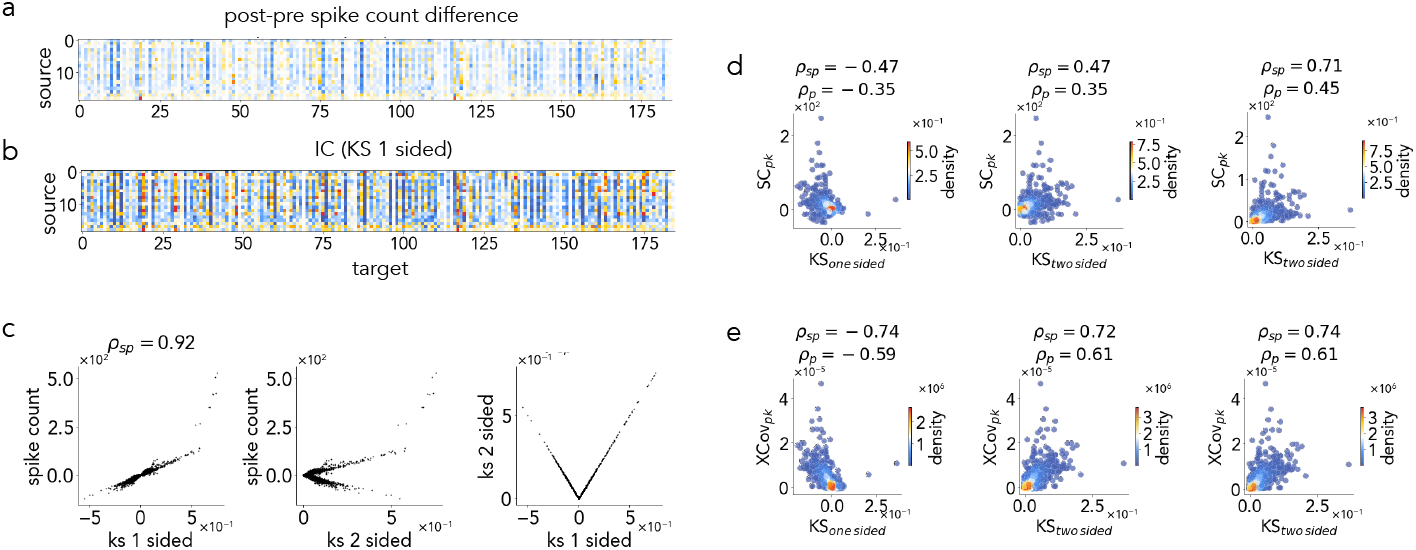
Signed IC. **(a)** Post–pre spike count differences computed using short windows (*T*_*w*_ = 50 ms) to isolate early response components. **(b)** Signed IC matrix obtained using the one-sided KS statistic. **(c)** Relationship between signed IC metrics. **Left**: spike-count difference vs. one-sided KS (Spearman *ρ* = 0.92). **Center**: spike-count difference vs. two-sided KS. **Right**: one-sided vs. two-sided KS. **(d)** Comparison between KS-based IC and signed/unsigned SC. Left: signed SC vs. one-sided KS (negative correlation). Middle: unsigned SC vs. one-sided KS. Right: unsigned SC vs. two-sided KS.

#### Note on the Markram-Tsodyks Model and the Depression-Only Limit

We summarize the classical Markram–Tsodyks (M–T) model [35, 36] and its reduction to the depression-only form used in our simulations. The model tracks three fractions of synaptic resources: *x*(*t*) (recovered/available), *y*(*t*) (active), and *z*(*t*) (inactive), with the conservation constraint *x*(*t*) + *y*(*t*) + *z*(*t*) = 1. For a presynaptic spike train {*t*_*i*_}, the governing equations are

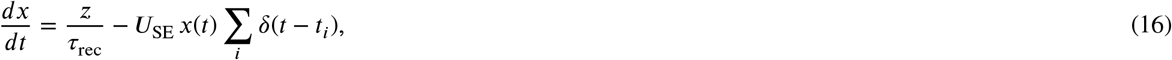

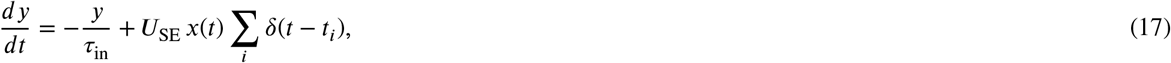

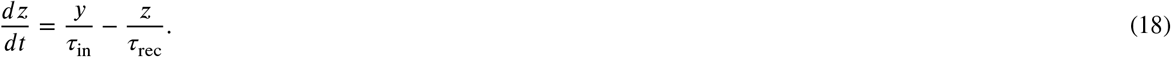

Here *U*_SE_ is the synaptic utilization, *τ*_rec_ is the recovery time constant, and *τ*_in_ is the inactivation time constant.

Summing Eqs. (16)-(18) yields

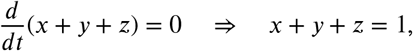

so *z*(*t*) can be eliminated, reducing the model to two variables (*x, y*). The equation for *y*(*t*) is of the form

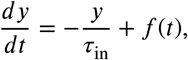

whose solution is

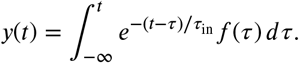

Thus *y*(*t*) acts as a fast low-pass filter of the spike-driven term *f* (*t*). In the limit of rapid inactivation (*τ*_in_ → 0),

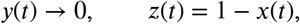

and the system reduces to the single equation

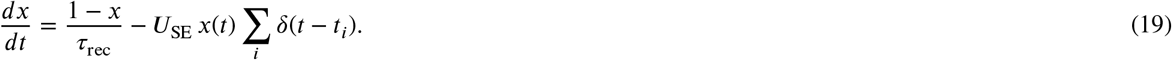

Between presynaptic spikes, Eq. (19) yields the recovery dynamics

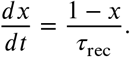

Integrating Eq. (19) across a spike at *t* = *t*_*i*_ gives the instantaneous update

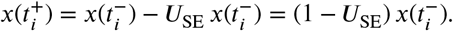

Defining *β* = 1 − *U*_SE_, this becomes

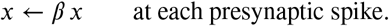

This is the depression-only formulation used in our spiking network, where the presynaptic resource variable *R*_*i*_(*t*) corresponds to *x*(*t*) in Eq. (19). This reduction is standard in the regime *τ*_in_ ≪ *τ*_rec_, consistent with fast vesicle activation followed by slow recovery observed in dissociated cultures.

## References

[1] C. D. Salzman, K. H. Britten, W. T. Newsome, Cortical microstimu-lation influences perceptual judgements of motion direction, Nature 346 (1990) 174–177.

[2] C. D. Salzman, C. M. Murasugi, K. H. Britten, W. T. Newsome, Microstimulation in visual area mt: Effects on direction discrimination performance, The Journal of Neuroscience 12 (1992) 2331–2355.

[3] C. M. Murasugi, C. D. Salzman, W. T. Newsome, Microstimulation in visual area mt: effects of varying pulse amplitude and frequency, The Journal of Neuroscience 13 (1993) 1719–1729.

[4] R. Romo, A. Hernández, A. Zainos, E. Salinas, Somatosensory discrimination based on cortical microstimulation, Nature 392 (1998) 387–390.

[5] M. S. Graziano, C. S. Taylor, T. Moore, Complex movements evoked by microstimulation of precentral cortex, Neuron 34 (2002) 841–851.

[6] D. J. O’Shea, L. Duncker, W. Goo, X. Sun, S. Vyas, E. M. Trautmann, I. Diester, C. Ramakrishnan, K. Deisseroth, M. Sahani, K. V. Shenoy, Direct neural perturbations reveal a dynamical mechanism for robust computation, bioRxiv (2022).

[7] A. Sanzeni, A. Palmigiano, T. H. Nguyen, J. Luo, J. J. Nassi, J. H. Reynolds, M. H. Histed, K. D. Miller, N. Brunel, Mechanisms underlying reshuffling of visual responses by optogenetic stimulation in mice and monkeys, Neuron 111 (2023) 1–17.

[8] K. Rakhshan, J. D. Schall, A. C. Huk, R. Kiani, Neural mechanisms underlying robust target selection in response to microstimulation of the oculomotor system, The Journal of Neuroscience 45 (2025) e2356232024.

[9] J. Soldado-Magraner, Y. Minai, B. M. Yu, M. A. Smith, Robustness of working memory to prefrontal cortex microstimulation, bioRxiv (2025).

[10] P. T. Sadtler, K. M. Quick, M. D. Golub, S. M. Chase, S. I. Ryu, E. C. Tyler-Kabara, B. M. Yu, A. P. Batista, Neural constraints on learning, Nature 512 (2014) 423–426.

[11] N. P. Shah, A. Phillips, S. Madugula, A. Sher, A. Litke, E. Chichilnisky, Precise control of neural activity using dynamically optimized electrical stimulation, eLife 13 (2024) RP83424.

[12] H. Y. Chau, K. D. Miller, Exact linear theory of perturbation response in a spaceand feature-dependent cortical circuit model, bioRxiv (2024).

[13] Y. Ghenbot, X. Liu, H. Hao, C. Rinehart, S. DeLuccia, S. T. Mal-donado, G. Boyek, M. Zhang, F. Aflatouni, J. Van der Spiegel, T. H. Lucas, A. G. Richardson, Goal-Directed BCI Feedback Using Cortical Microstimulation, Springer International Publishing, Cham, 2020, pp. 65–73. URL: https://doi.org/10.1007/978-3-030-49583-1_7. doi:10.1007/978-3-030-49583-1_7.

[14] N. D. Shelchkova, J. E. Downey, C. M. Greenspon, E. V. Okorokova, R. Sobinov, C. Verbaarschot, Q. He, C. Sponheim, A. F. Tortolani, D. D. Moore, M. T. Kaufman, R. C. Lee, D. Satzer, J. Gonzalez-Martinez, P. C. Warnke, L. E. Miller, M. L. Boninger, R. A. Gaunt, J. L. Collinger, N. G. Hatsopoulos, S. J. Bensmaia, Microstimulation of human somatosensory cortex evokes task-dependent, spatially patterned responses in motor cortex, Nature Communications 14 (2023).

[15] R. Viaro, D. Bernardi, E. Maggiolini, A. D’Ausilio, C. G. Ferroni, P. Parmiani, L. Fadiga, Differential motor neuron activity in rats during successful and failed grasping, Cerebral Cortex 35 (2025) bhaf032.

[16] G. Barzon, A. De, I. Moran, C. Carnahan, L. Mazzucato, R. Kiani, Control of cortical population activity with patterned microstimulation, bioRxiv (2026).

[17] U. Topalovic, S. Barclay, C. Ling, A. Alzuhair, W. Yu, V. Hokhikyan, H. Chandrakumar, D. Rozgic, W. Jiang, S. Basir-Kazeruni, S. L. Maoz, C. S. Inman, A. Bari, A. Fallah, D. Eliashiv, I. Fried, N. Suthana, D. Markovic, A wearable platform for closed-loop stimulation and recording of single-neuron and local field potential activity in freely moving humans, Nature Neuroscience (2023).

[18] C. R. Oehrn, S. Cernera, L. H. Hammer, et al., Chronic adaptive deep brain stimulation versus conventional stimulation in parkinson’s disease: a blinded randomized feasibility trial, Nature Medicine 30 (2024) 3345–3356.

[19] R. Fisher, A. L. Velasco, Electrical brain stimulation for epilepsy, Nature Reviews Neurology 10 (2014) 261–270.

[20] E. Cohen, M. Ivenshitz, V. Amor-Baroukh, V. Greenberger, M. Se-gal, Determinants of spontaneous activity in networks of cultured hippocampus, Brain Research 1235 (2008) 21–30.

[21] J. Soriano, M. R. Martínez, T. Tlusty, E. Moses, Development of input connections in neural cultures, Proceedings of the National Academy of Sciences of the United States of America 105 (2008) 13758–13763.

[22] M. Chiappalone, M. Bove, A. Vato, M. Tedesco, S. Martinoia, Dis-sociated cortical networks show spontaneously correlated activity patterns during in vitro development, Brain Research 1093 (2006) 41–53.

[23] D. A. Wagenaar, J. Pine, S. M. Potter, An extremely rich repertoire of bursting patterns during the development of cortical cultures, BMC Neuroscience 7 (2006) 11.

[24] A. Mazzoni, F. D. Broccard, E. Garcia-Perez, P. Bonifazi, M. E. Ruaro, V. Torre, On the dynamics of the spontaneous activity in neuronal networks, PLoS ONE 2 (2007) e439.

[25] V. Pasquale, P. Massobrio, L. L. Bologna, M. Chiappalone, S. Martinoia, Self-organization and neuronal avalanches in networks of dissociated cortical neurons, Neuroscience 153 (2008) 1354–1369.

[26] S. Okujeni, S. Kandler, U. Egert, Mesoscale architecture shapes initiation and richness of spontaneous network activity, Journal of Neuroscience 37 (2017) 3972–3987.

[27] I. Colombi, T. Nieus, M. Massimini, M. Chiappalone, Spontaneous and perturbational complexity in cortical cultures, Brain Sciences 11 (2021).

[28] M. E. Obien, K. Deligkaris, T. J. Bullmann, D. J. Bakkum, U. Frey, Revealing neuronal function through microelectrode array recordings, Frontiers in Neuroscience 8 (2015) 423.

[29] S. Ronchi, M. Fiscella, C. Marchetti, V. Viswam, J. Müller, U. Frey, Hierlemann, Single-cell electrical stimulation using cmos-based high-density microelectrode arrays, Frontiers in neuroscience 13 (2019) 208.

[30] A. Nejatbakhsh, F. Fumarola, S. Esteki, T. Toyoizumi, R. Kiani, L. Mazzucato, Predicting the effect of micro-stimulation on macaque prefrontal activity based on spontaneous circuit dynamics, Phys. Rev. Res. 5 (2023) 043211.

[31] S. Sadeh, C. Clopath, Theory of neuronal perturbome in cortical networks, Proceedings of the National Academy of Sciences 117 (2020) 26966–26976.

[32] N. Wiener, The theory of prediction, in: E. Beckenbach (Ed.), Modern Mathematics for Engineers, McGraw-Hill, New York, 1956.

[33] S. Ito, M. Hansen, R. Heiland, A. Lumsdaine, A. Litke, J. Beggs, Extending transfer entropy improves identification of effective connectivity in a spiking cortical network model, in: PLoS One, 2011.

[34] P. Antonello, T. Varley, J. Beggs, M. Porcionatto, O. Sporns, J. Faber, Self-organization of in vitro neuronal assemblies drives to complex network topology, Elife (2022).

[35] M. Tsodyks, H. Markram, The neural code between neocortical pyramidal neurons depends on neurotransmitter release probability, Proceedings of the National Academy of Sciences 94 (1997) 719–723.

[36] M. Tsodyks, K. Pawelzik, H. Markram, Neural networks with dynamic synapses, Neural Computation 10 (1998) 821–835.

[37] E. Alvarez-Lacalle, E. Moses, Slow and fast pulses in 1-d cultures of excitatory neurons, Journal of Computational Neuroscience 26 (2009) 475–493.

[38] H. Yamamoto, F. P. Spitzner, T. Takemuro, V. Buendía, H. Murota, C. Morante, T. Konno, S. Sato, A. Hirano-Iwata, A. Levina, V. Priesemann, M. A. Muñoz, J. Zierenberg, J. Soriano, Modular architecture facilitates noise-driven control of synchrony in neuronal networks, Science Advances 9 (2023) eade1755.

[39] M. Montalà-Flaquer, C.F. López-León, D. Tornero, A. M. Houben, T. Fardet, P. Monceau, S. Bottani, J. Soriano, Rich dynamics and functional organization on topographically designed neuronal networks in vitro, Iscience 25 (2022).

[40] Y. Lu, Z. Wang, X. Li, Y. Wang, Y. Wang, X. Zhao, J. Zhang, J. Luo, Unveiling the impact of low-frequency electrical stimulation on network synchronization and learning behavior in cultured hippocampal neural networks, Brain Research 1810 (2024) 148163.

[41] T. Kobayashi, K. Shimba, T. Narumi, T. Asahina, K. Kotani, Y. Jimbo, Revealing single-neuron and network-activity interaction by combining high-density microelectrode array and optogenetics, Nature Communications 15 (2024) 9547.

[42] A. M. Houben, J. Garcia-Ojalvo, J. Soriano, Role of connectivity anisotropies in the dynamics of cultured neuronal networks, arXiv preprint 2501.04427 (2025).

[43] B. J. Kagan, A. C. Kitchen, N. T. Tran, F. Habibollahi, M. Khajehne-jad, B. J. Parker, A. Bhat, B. Rollo, A. Razi, K. J. Friston, In vitro neurons learn and exhibit sentience when embodied in a simulated game-world, Neuron 110 (2022) 3952–3969.

[44] T. Sumi, H. Yamamoto, Y. Katori, K. Ito, S. Moriya, T. Konno, S. Sato, A. Hirano-Iwata, Biological neurons act as generalization filters in reservoir computing, Proceedings of the National Academy of Sciences 120 (2023) e2217008120.

[45] F. Borra, S. Cocco, R. Monasson, Task learning through stimulation-induced plasticity in neural networks, PRX Life 2 (2024) 043014.

[46] S. Hua, Y. Liu, J. Luo, S. Li, L. Jiang, P. Wu, S. Sun, L. Shang, C. Lu, K. Zhang, et al., Microelectrode arrays cultured with in vitro neural networks for motion control tasks: encoding and decoding progress and advances, Microsystems & Nanoengineering 11 (2025) 233.

[47] A. Yaron, Z. Zhang, D. Akita, T. I. Shiramatsu, Z. C. Chao, H. Takahashi, Dissociated neuronal cultures as model systems for self-organized prediction, Frontiers in Neural Circuits 19 (2025) 1568652.

[48] C. Beer, O. Barak, Revealing and reshaping attractor dynamics in large networks of cortical neurons, PLOS Computational Biology 20 (2024) e1011784.

[49] A. Ahtiainen, L. Leydolph, J. M. A. Tanskanen, A. Hunold, J. Haueisen, J. A. K. Hyttinen, Electric field temporal interference stimulation of neurons in vitro, Lab Chip 24 (2024) 3945–3957.

[50] J. Wen, M. Peitz, O. Brüstle, A defined human-specific platform for modeling neuronal network stimulation in vitro and in silico, Journal of Neuroscience Methods 373 (2022) 109562.

[51] N. R. Wilson, F. L. Wang, N. Chen, S. X. Yan, A. L. Daitch, Shi, S. Sharma, M. Sur, A platform for spatiotemporal “matrix” stimulation in brain networks reveals novel forms of circuit plasticity, Frontiers in Neural Circuits 15 (2022) 792228.

[52] S. N. Chettih, C. D. Harvey, Single-neuron perturbations reveal feature-specific competition in v1, Nature 567 (2019) 334–340.

[53] A. M. Packer, B. Roska, M. Häusser, Targeting neurons and photons for optogenetics, Nature neuroscience 16 (2013) 805–815.

[54] N. Seseri, D. Nigrisoli, F. Faraci, A. D’Angelo, R. Freddi, S. Chandran, R. Barbieri, S. Corti, L. Ottoboni, S. Russo, Electrical stimulation elicits space-and parameter-dependent spiking responses in human cortical organoids, bioRxiv (2025) 2025–09.

[55] T. Sharf, T. Van Der Molen, S. M. Glasauer, E. Guzman, A. P. Buccino, G. Luna, Z. Cheng, M. Audouard, K. G. Ranasinghe, K. Kudo, et al., Functional neuronal circuitry and oscillatory dynamics in human brain organoids, Nature communications 13 (2022) 4403.

[56] T. Van der Molen, Studying spontaneous and evoked neural circuit activity using brain organoids, UC Santa Barbara, 2025.

[57] V. Pasquale, S. Martinoia, M. Chiappalone, Stimulation triggers endogenous activity patterns in cultured cortical networks, Scientific reports 7 (2017) 9080.

[58] T. Opitz, A. D. De Lima, T. Voigt, Spontaneous development of synchronous oscillatory activity during maturation of cortical networks in vitro, Journal of neurophysiology 88 (2002) 2196–2206.

[59] P. Schnepel, A. Kumar, M. Zohar, A. Aertsen, C. Boucsein, Physiol-ogy and impact of horizontal connections in rat neocortex., Cerebral Cortex 25 (2014) 3818–3835.

[60] S. Horvát, R. Gămănut, M. Ercsey-Ravasz, L. Magrou, B. Gămănut, C. Van Essen, A. Burkhalter, K. Knoblauch, Z. Toroczkai, H. Kennedy, Spatial embedding and wiring cost constrain the functional layout of the cortical network of rodents and primates, PLoS biology 14 (2016) e1002512.

[61] Z.-S. Lv, C.-P. Zhu, P. Nie, J. Zhao, H.-J. Yang, Y.-J. Wang, C.-K. Hu, Exponential distance distribution of connected neurons in simulations of two-dimensional in vitro neural network development, Frontiers of Physics 12 (2017) 128902.

[62] P. Verstraelen, M. Van Dyck, M. Verschuuren, N. D. Kashikar, R. Nuydens, J.-P. Timmermans, W. H. De Vos, Image-based profiling of synaptic connectivity in primary neuronal cell culture, Frontiers in Neuroscience 12 (2018) 389.

[63] P. Verstraelen, M. Verschuuren, W. H. De Vos, Integrated staging of morphofunctional connectivity in neuronal cultures, STAR protocols 5 (2024) 102957.

[64] J. T. Savage, J. J. Ramirez, W. C. Risher, Y. Wang, D. Irala, C. Eroglu, Synbot is an open-source image analysis software for automated quantification of synapses, Cell Reports Methods 4 (2024).

[65] T. M. Cramer, S. K. Tyagarajan, Protocol for the culturing of primary hippocampal mouse neurons for functional in vitro studies, STAR protocols 5 (2024) 102991.

[66] D. Bernardi, B. Lindner, Detecting single-cell stimulation in a large network of integrate-and-fire neurons, Physical Review E 99 (2019) 032304.

[67] D. Bernardi, G. Doron, M. Brecht, B. Lindner, A network model of the barrel cortex combined with a differentiator detector reproduces features of the behavioral response to single-neuron stimulation, PLOS Computational Biology 17 (2021) 1–38.

[68] A. A. Tsonis, E. R. Deyle, H. Ye, G. Sugihara, Convergent cross mapping: theory and an example, Advances in nonlinear geosciences (2017) 587–600.

[69] M. Maschietto, S. Girardi, M. Dal Maschio, M. Scorzeto, S. Vas-sanelli, Sodium channel β 2 subunit promotes filopodia-like processes and expansion of the dendritic tree in developing rat hippocampal neurons, Frontiers in Cellular Neuroscience 7 (2013).

[70] D. A. Wagenaar, J. Pine, S. M. Potter, Effective parameters for stimu-lation of dissociated cultures using multi-electrode arrays, Journal of neuroscience methods 138 (2004) 27–37.

[71] I. E. M., Simple model of spiking neurons, IEEE Transactions on Neural Networks 14 (2003) 1569–1572.

[72] I. EM, Polychronization: computation with spikes, Neural Computa-tion 18 (2006) 245–282.

[73] P. Dayan, L. F. Abbott, Theoretical Neuroscience: Computational and Mathematical Modeling of Neural Systems, MIT Press, Cambridge, MA, 2001.

[74] D. Golomb, Y. Amitai, Propagating neuronal discharges in neocortical slices: computational and experimental study, Journal of Neurophysiology 78 (1997) 1199–1211.

[75] S. Butovas, S. G. Hormuzdi, H. Monyer, C. Schwarz, Effects of electrically coupled inhibitory networks on local neuronal responses to intracortical microstimulation., Journal of Neurophysiology 96 (2006) 1227–1236.

[76] P. E. Latham, B. J. Richmond, P. G. Nelson, S. Nirenberg, Intrinsic dynamics in neuronal networks. i. theory, Journal of Neurophysiology 83 (2000) 808–827. PMID: 10669496.

[77] P. E. Latham, B. J. Richmond, S. Nirenberg, P. G. Nelson, Intrinsic dynamics in neuronal networks. ii. experiment, Journal of Neurophysiology 83 (2000) 828–835. PMID: 10669497.

[78] A. Bucci, M. Büttner, N. Domdei, F. B. Rosselli, M. Znidaric, R. Diggelmann, M. De Gennaro, C. S. Cowan, W. Harmening, A. Hierlemann, B. Roska, F. Franke, Action potential propagation speed compensates for traveling distance in the human retina, bioRxiv preprint (2024).

[79] T. Baltz, T. Voigt, Interaction of electrically evoked activity with intrinsic dynamics of cultured cortical networks with and without functional fast gabaergic synaptic transmission, Frontiers in Cellular Neuroscience Volume 9 - 2015 (2015).

[80] M. Brecht, M. Schneider, B. Sakmann, T. W. Margrie, Whisker movements evoked by stimulation of single pyramidal cells in rat motor cortex., Nature 427 (2004) 704–710.

[81] D. Bernardi, B. Lindner, Optimal detection of a localized perturbation in random networks of integrate-and-fire neurons, Physical Review Letters 118 (2017) 268301.

[82] A. R. Houweling, M. Brecht, Behavioural report of single neuron stimulation in somatosensory cortex., Nature 451 (2008) 65–68.

[83] V. Pastore, P. Massobrio, A. Godjoski, S. Martinoia, Identification of excitatory-inhibitory links and network topology in large-scale neuronal assemblies from multi-electrode recordings, PLOS Computational Biology 14 (2018) e1006381.

